# Multivalency in endocytic accessory proteins orchestrates membrane shape transitions

**DOI:** 10.64898/2026.10.26.762025

**Authors:** Honey Priya James, Akshatha Manjunatha, Olivia Bullock, Tobias Baumgart

**Affiliations:** University of Pennsylvania

## Abstract

Fast endophilin-mediated endocytosis (FEME) relies on the coordinated activities of membrane-remodeling proteins and multivalent protein interaction networks. Endophilin is a potent membrane curvature generator through its N-BAR domain and forms multivalent interactions with the proline-rich motifs (PRMs) of lamellipodin via its SH3 domain. Although these interactions are known to promote biomolecular condensation, their functional consequences for membrane remodeling have remained unclear. Here, we investigate how SH3/PRM-mediated phase separation regulates endophilin-driven membrane curvature generation. Using purified recombinant proteins, giant unilamellar vesicles, quantitative fluorescence microscopy, and in vitro phase-separation assays, we show that multivalent interactions between endophilin and lamellipodin drive the formation of membrane-associated protein-rich phases that strongly suppress endophilin-mediated membrane tubulation. Competitive disruption of SH3/PRM interactions with the synaptojanin-derived peptide PP19 (containing only two PRM motifs) rapidly dissolves condensates and restores membrane curvature generation without affecting the intrinsic curvature-generating activity of endophilin. We further demonstrate that this regulatory mechanism is conserved across endophilin A1, A2, and A3 isoforms and depends critically on interaction valency: a multivalent PRM7 construct recapitulates condensate formation and curvature suppression, whereas mono- or divalent PRM constructs does not. Together, these findings identify multivalent SH3/PRM interactions as a reversible molecular switch that balances biomolecular condensation and membrane remodeling. Multiple mechanisms, including post-translational modifications, likely regulate the multivalency of these interactions. Our results thus establish a broad functional link between protein phase separation and membrane mechanics, providing a framework for understanding how multivalent interaction networks regulate membrane shaping in multiple endocytic pathways.

## Introduction

Membrane remodeling is fundamental to endocytosis, intracellular trafficking and signal transduction. These processes require coordinated changes in membrane curvature together with the recruitment and organization of protein interactions that control when and where the deformation occurs[1–5]. Curvature generating proteins therefore do not act in isolation but function within dense assemblies of adaptors, scaffolds and signaling proteins[6, 7]. Understanding how these interactions influence the membrane remodeling proteins is essential for explaining how membrane shape is dynamically regulated in cells. Fast endophilin mediated endocytosis (FEME) is a clathrin independent endocytic pathway that depends critically on endophilin family proteins and their interaction partners[8, 9]. Several additional endocytic pathways are endophilin dependent, including ultrafast endocytosis[10–14]. Endophilin combines two distinct functions within a single molecule. Its N BAR domain binds membranes and the protein dimer generates curvature through scaffolding and oligomerization, whereas its C-terminal SH3 binding domain interacts with proline rich motifs (PRMs) present in endocytic accessory proteins[15, 16]. This domain organization places endophilin at an important regulatory interface between membrane mechanics and multivalent protein interaction networks.

Recent studies have revealed that many endocytic proteins form biomolecular condensates through liquid-liquid phase separation (LLPS), driven by multivalent interactions among modular binding domains and intrinsically disordered regions [17–22]. Such condensates can concentrate signaling molecules, organize reaction networks, and generate spatially distinct membrane-associated protein phases [23, 24]. Endophilin itself has been shown to undergo phase separation through self-association of its BAR domain [25], while multivalent interactions between its SH3 domain and proline-rich motifs (PRMs) of binding partners such as lamellipodin further promote condensate formation [25, 26]. Moreover, membrane-associated clustering of endophilin and lamellipodin has been proposed to contribute to the initiation and regulation of FEME [25, 27, 28]. However, the effect of lamellipodin-driven multivalent assembly on the intrinsic membrane-remodeling activity of endophilin has not yet been fully resolved. Although the molecular basis of endophilin-LPD phase separation has begun to emerge, its functional consequences for membrane remodeling remain poorly understood. Endophilin is a potent generator of membrane curvature [15, 29], whereas the formation of dense multivalent protein assemblies could alter protein mobility, organization, and access to the membrane [18, 23]. Previous reconstitution experiments showed that lamellipodin binding causes shortening and disappearance of endophilin-generated membrane tubules [25]. This observation suggests that multivalent assembly may negatively regulate membrane curvature generation, but several mechanistic questions remain unanswered. It is unclear whether curvature suppression is directly linked to the formation of multivalent endophilin-lamellipodin assemblies, whether the inhibited state can be reversed after it has formed, and how strongly this regulation depends on interaction valency. In particular, how SH3-PRM interaction valency and competitive binding regulate the balance between condensation and membrane tubulation remains unresolved. In addition, because endophilin SH3 domains interact with multiple PRM containing proteins, competition among binding partners could provide a mechanism for dynamically reorganizing these assemblies. The synaptojanin derived peptide-PP19 which contains two PRM motifs and binds endophilin SH3 domains provides a useful model for testing whether competitive redistribution of SH3-PRM interactions can reverse LPD-dependent assembly and restore membrane remodeling activity. Furthermore, vertebrates express three closely related endophilin A isoforms (A1, A2, and A3) [30], yet it remains unclear whether condensate formation and its effects on membrane remodeling are conserved across the family.

Here, we investigate how multivalent interactions between endophilin and lamellipodin regulate membrane curvature generation and phase separation. Using giant unilamellar vesicles, purified proteins, and quantitative fluorescence microscopy, we show that multivalent SH3-PRM interactions drive the formation of membrane-associated protein-rich phases that suppress endophilin-mediated membrane tubulation. Disruption of these interactions by PP19 [31] dissolves condensates and restores membrane curvature generation. We further demonstrate that this regulatory mechanism is conserved across endophilin A isoforms and depends critically on PRM valency. Together, our findings imply that multivalent protein interactions, whether orchestrated by competitive binding interactions or post-translational modifications, can function as a switch that balances phase separation and membrane remodeling, providing a new framework for understanding the regulation of endophilin-dependent endocytic processes.

## Materials and Methods

### Materials

Tris hydrochloride (Tris-HCL), Tris 2-carboxyethyl Phosphine hydrochloride (TCEP-HCL), (4-2-(hydroxyethyl)-1-Piperazineethanesulfonic acid (HEPES), Sodium Chloride (NaCl), Ethylenediaminetetraacetic acid (EDTA), Dithiothreitol (DTT), Phenylmethanesulfonylfluoride (PMSF), Isopropyl-β-D-thiogalactopyranoside (IPTG), Glutathione (GSH), Sodium dihydrogenphosphate (NaH2PO4), Disodium hydrogen phosphate (Na2HPO4), Imidazole, Sodium hydroxide (NaOH), Sucrose, and Chloroform were procured from Fisher Scientific. β-Casein was obtained from Sigma Aldrich. Indium Tin Oxide (ITO) slides were obtained from Delta Technologies (CG-40IN-S115). Custom-synthesized PP19 (Ac-CVAPPARPAPPQRPPPPSGA-NH2) was purchased from (Thermo Fisher Scientific), LPD C4 (Ac-KKPPPTPQRNSSI-NH2) was obtained from Peptide 2.0, Alexa fluor 647^TM^-C2 maleimide, Alexa fluor 488^TM^-C5 maleimide, and Alexa fluor 647^TM^-NHS ester (Succinimidyl ester) were procured from Invitrogen. Lipids 1,2-dioleolyl-sn-glycero-3-phosphocholine (DOPC), 1,2-dioleolyl-sn-glycero-3-phosphoethanolamine (DOPE), and 1,2-dioleolyl-sn-glycero-3-phospho-L-Serine (DOPS) in chloroform were purchased from Avanti Polar Lipids Inc. (Al, USA). 1,2-Dihexadecanoyl-sn-Glycero-3-phosphoethanolamine (Marina Blue DHPE) was procured from Invitrogen. All reagents were used without further purification.

### Plasmids

Rat-endophilin A1(with C108S, E241C, C294S, C295S mutations) was described elsewhere [32]. Sequences encoding rat-endophilin A2, rat-endophilin A3, and LPD 850-1250 were synthesized and cloned into a pGEX-6P1 vector by Biomatik, USA. The Plasmid encoding seven repeats of PRM from lamellipodin (aa 970-981, FHMCGGSWGGSGGSGGSKKPPPTPQRNSSIGGSGGSGGSGGSKKPPPTPQRNSSIGGSGG SGGSGGSKKPPPTPQRNSSIGGSGGSGGSGGSKKPPPTPQRNSSIGGSGGSGGSGGSKKPP PTPQRNSSIGGSGGSGGSGGSKKPPPTPQRNSSIGGSGGSGGSGGSKKPPPTPQRNSSIGSR),

containing (Gly-Gly-Ser)4 linkers and an N-terminal tryptophan, was custom-synthesized by Biomatik, USA. The plasmid pMAL-Abl-PRM 5R (Addgene plasmid 112088) was a generous gift from Michael Rosen’s lab. The PRM7 sequence was cloned into pMAL-Abl-PRM 5R by replacing its PRM 5 sequence. The TEV cleavage site between PRM-7 and the C-terminal 6x his tag was mutated to prevent the removal of the His tag during purification.

### Protein expression and purification

BL21DE3 Codon Plus (RIL) cells (Agilent technologies) were transformed with plasmids encoding rat Endophilin A1, A2, A3 and LPD 850-1250 (ampicillin resistant) or PRM-7 (Kanamycin resistant) and grown on LB Agar plates containing respective antibiotics (ampicillin 100 µg/ml, kanamycin 50 µg/ml). Single colonies from the transformed plates were selected to inoculate the overnight primary culture (100 ml of LB media supplemented with the corresponding antibiotic) shaken at 250 rpm at 37°C. Larger volume cultures (4 ×1L for EDP A1, EDP A2, EDP A3, LPD 850-1250 and PRM 7) were grown using a 25 ml of primary culture per liters of LB media supplemented with same antibiotic, at 37°C, shaken at 225 rpm until the optical density (O.D) at 600 nm reached 0.6 – 0.7. The cultures were cooled and induced at 18°C with IPTG (0.6 mM for EDPA1, EDPA2, EDPA3, PRM7 and 0.3 mM for LPD 850-1250) for 16-18 h. Cells were harvested by spinning down at 6000 rpm for 10 min at 4°C, and the pellet was resuspended in a GST lysis buffer (300 mM NaCl, 50 mM Tris, 1mM EDTA, 2 mM DTT, 1mM PMSF, pH 8) for endophilin A1, A2 and A3. The aforementioned lysis buffer additionally contained 20 mM imidazole and no DTT nor EDTA (Ni-NTA lysis buffer) for LPD 850-120 or PRM 7. The bacterial cell suspension was lysed using tip sonication (Qsonica Q700) and centrifuged at 14000 rpm for 1 h to remove the cellular debris. The supernatant was filtered through 0.22 µm syringe filter (Millipore) and further purified by fast protein liquid chromatography (FPLC, AKTA Pure). SDS-PAGE was performed between each chromatography step to assess the purity of the protein. The protein concentrations were determined by measuring the absorbance at 280 nm using a nanodrop spectrophotometer (Nanodrop 2000c). Endophilin concentrations are displayed as monomer concentrations throughout. Further purification specific details for each protein are mentioned below.

### Endophilin A1, A2 and A3

The endophilin isoforms (A1, A2 and A3) were extracted from the respective filtered supernatant with 3 GSTrap FF columns (5ml, Cytiva), connected in series, and pre-equilibrated with GST lysis buffer. The columns were washed extensively with GST wash buffer (150 mM NaCl, 50 mM Tris, 1mM EDTA, 2 mM DTT, pH 8) followed by elution of GST-tagged proteins with elution buffer (150 mM NaCl, 50 mM Tris, 20 mM GSH, 1 mM EDTA, 2 mM DTT, pH 8). The eluted protein was incubated with PreScission protease at 4°C overnight to remove the GST tag. The cleaved mixture was then loaded onto an anion exchange column (Hi Trap Q FF, 5 ml, Cytiva) equilibrated with anion exchange buffer QA (150 mM NaCl, 50 mM Tris, 1 mM EDTA, 1 mM DTT, pH 8). The Q column was then washed with QA buffer, and the proteins were then eluted with a linear gradient of 0 - 75% anion exchange buffer QB (1M NaCl, 50 mM Tris, 1 mM EDTA, 1 mM DTT, pH 8), and the peak fractions were analyzed by SDS-PAGE. The fractions containing the proteins were pooled, concentrated and loaded onto a size exclusion column (Superdex 200 pg 16/600 Hi Load, Cytiva) equilibrated with storage buffer (150 mM NaCl, 20 mM HEPES, 1 mM TCEP, pH 7.4). The pure fractions were pooled and concentrated by spin centrifugation, flash frozen in liquid nitrogen and stored at -80°C for further use.

### LPD 850-1250

LPD 850-1250 (referred to as LPD in the main text) was expressed with an N-terminal GST tag and C-terminal His-tag. The filtered supernatant containing LPD 850-1250 was incubated overnight with Ni-NTA resin (HisPur, Thermoscientific) equilibrated with Ni-NTA lysis buffer at 4°C under gentle agitation. The protein-bound resin was washed with Ni-NTA lysis buffer and incubated overnight at 4°C with precision protease to cleave the GST tag. The resin was further washed with Ni-NTA lysis buffer, and the protein was eluted with high imidazole-containing Ni-NTA elution buffer (150 mM NaCl, 50 mM Tris, 500 mM Imidazole, pH ∼8). The eluted protein was buffer exchanged to cation exchange buffer SP A (150 mM NaCl, 20 mM sodium phosphate, 1 mM EDTA, 1mM DTT, pH ∼7) using desalting column (HiTrap, 5 ml, Cytiva). The buffer-exchanged protein was loaded onto a cation exchange column (HiTrap SP HP, 5ml, Cytiva) equilibrated with SP A buffer. The column was then washed with SP A buffer, and the bound protein was then eluted with a linear gradient of 0-45% cation exchange buffer SP B (1 M NaCl, 20 mM sodium phosphate, 1 mM EDTA, 1mM DTT, pH ∼7). The protein containing fractions were then pooled, concentrated and further purified by size exclusion chromatography using size exclusion column (Superdex 200 pg 16/600 Hi Load, Cytiva). The pure fractions are concentrated, flash frozen in liquid nitrogen and stored at -80°C.

### PRM7

The PRM-7 his construct was expressed as an N-terminal MBP and C-terminal his tag protein. The filtered supernatant was loaded onto an HisTrap HP affinity column (5ml, Cytiva) equilibrated with Ni-NTA lysis buffer. After washing the bound protein with lysis buffer, elution was performed using Ni-NTA elution buffer. The eluted sample was loaded onto a HiTrap SP HP cation exchange column, and the protein was eluted with a linear gradient using SP A buffer and SP B buffer. The PRM-7 His fraction was dialysed at 4°C for 12 h against 2×1l of QA buffer using a dialysis membrane of 3.5 kDa molecular weight cut off (MWCO). The dialysed protein was applied to a HiTrap Q HP anion exchange column and MBP-PRM7-His eluted as flow through. The flow through fractions was incubated with TEV protease at 4°C for 12 h and following the cleavage the mixture was applied to amylose resin (New England Biolabs). The purified PRM-7 His protein was collected in the flow through, concentrated and exchanged to storage buffer. The proteins were flash frozen in liquid nitrogen and stored at -80°C.

### Fluorophore conjugation of proteins

Endophilin A1 and PRM-7 were labelled with Alexa Fluor-488 C5 maleimide (AF-488) and Alexa Fluor-647 C2 maleimide (AF-647) respectively. The fluorophore was added to the protein at a five-fold molar excess and allowed to react for 12-16 h. LPD 850-1250 was labelled at the N -terminus with Alexa fluor 647-Succinimidyl ester. Before N-terminus labelling, the protein was exchanged to an amine-free buffer (20 mM Sodium phosphate, 150 mM NaCl, pH∼7) and the fluorophore was added to the protein in five-fold molar excess and incubated for 12 h at 4°C. Labelled proteins were purified from unreacted dye using a desalting column (HiTrap, 5 ml, Cytiva). The final protein concentration after labelling was determined from the absorbance measurements at 280 nm via nanodrop. For Alexa -fluor 488, protein concentration (M) was calculated as (*A*_280_ − (*A*_494_ × 0.11))⁄*εε_Protein_* ; for Alexa-fluor 647 as (*A*_280_ − (*A*_650_ × 0.03))⁄*ε_Protein_* , where *A* represents the absorbance at the specified wavelength, 0.11 and 0.03 are the correction factors for AF-488 and AF-647 respectively, *ε_Protein_* is the extinction coefficient of the protein.

### Preparation of Giant Unilamellar Vesicles (GUV)s

GUVs of lipid composition DOPC: DOPS: DOPE: MB-DHPE (24.5:45:30:0.5 mole percent) were prepared using the electroformation method as described elsewhere[33]. Briefly, 10 µl of lipid mixture (5 mg/ml) in chloroform was spread into a thin lipid film onto conductive side of two indium tin oxide (ITO) slides and kept under vacuum for 2h to ensure the complete removal of chloroform. The lipid film was gently hydrated with 450 µl of 300 mM sucrose solution, after which the two-ITO coated slides were placed with their conductive sides facing each other, separated by a silicone spacer of thickness 2 mm and clipped together to assemble the chamber. An alternating current of amplitude 4 volts peak-to-peak and frequency 4 Hz was applied across two ITO slides for 2 h at room temperature using a function generator (Rigol, DG1000Z). Thereafter, the GUVs were carefully collected and stored in microcentrifuge tube (2 ml) and allowed to rest for 12 h before further use.

For the GUVs tubulation assay, the desired concentration of proteins and PP19 was diluted into an osmotically balanced experimental buffer (150 mM NaCl, 20 mM HEPES, 1mM TCEP, pH 7.4) matched to the osmolarity of the sucrose solution used during GUV formation. Subsequently, 2 µl of the GUV suspension was added to each reaction to obtain a final assay volume of about 10 µl. Imaging was performed in a closed chamber assembled by sandwiching samples between two 25 mm glass coverslips (Fisher brand, 12542C) separated by a thin layer of grease (Dow Corning) around the edge. The coverslips were passivated with 5 µl of 2mg/ml β-Casein in Milli Q water.

### Protein condensate experiments

The condensates composed of endophilin-LPD 850-1250 or endophilin-PRM 7 were formed by mixing the specified protein concentrations (see Fig 3B and Fig 7A) in the experimental buffer and incubated for 10 min at room temperature. The condensates were deposited on PEG-PLL passivated coverslips[34] and imaged using confocal microscopy. For the PP19 concentration-dependent study, the PP19 (0.5 to 100 µM) was added to the pre-formed condensates and incubated for 15 min at room temperature before imaging. For the condensate experiments, the coverslips (25 mm, Fisher scientific) were cleaned by bath sonication in 2% Helmanex, followed by three rinses with water. The cleaned coverslips were either stored immediately or stored in ethanol until use. Coverslips were passivated by coating with PEG-PLL as described previously[35]. The size of the condensates was measured using Fiji Image J.

### Confocal microscopy

Confocal imaging experiments were performed on an Olympus IX83 inverted microscope (Olympus, Centre Valley, PA) equipped with 60x / 1.2 numerical aperture water immersion objective. Sequential line scanning acquisition was used to avoid cross talk between the channels for multifluorophore samples. Excitation was achieved with lasers 405, 488, and 647 for Marina Blue DHPE, Alexa fluor-488 and Alexa fluor 647 respectively.

### Statistical data analysis of membrane tubulation

Outward tubulation was defined as the presence of outer membrane tubes, either alone or together with inner tubes. Tubulation frequencies were analyzed using binomial logistic regression, with treatment concentration as a categorical fixed effect and experimental trial as a blocking factor. Extra-binomial variability was accounted for by adjusting coefficient variances using the Pearson dispersion factor. Overall treatment effects were assessed using Wald F-tests. Treatment–control comparisons were performed using two-sided Wald tests with Dunnett adjustment for multiple comparisons, whereas individual treatment/control comparisons were evaluated using two-sided Wald tests without multiplicity adjustment.

## Results

### Lamellipodin 850-1250 and Endophilin A1 independently drive membrane curvature generation

To investigate the role of multivalent SH3-PRM interactions in regulating endophilin mediated membrane curvature, we first examined the curvature-generation capacities of endophilin and Lamellipodin independently using a GUV-based tubulation assay. Lamellipodin (amino acid residues 1-1250) contains coiled-coil (CC), Ras-associated (RA) and Pleckstrin homology (PH) domains followed by an intrinsically disordered C-terminal region containing proline-rich motifs (PRMs) (Figure 1A). The disordered region (658 amino acids) contains ten PRMs which have been identified in previous studies as SH3 binding sites of endophilin[36]. To facilitate protein expression and purification[37], we utilized a truncated construct (residues 850-1250) of lamellipodin hereafter referred as LPD. This segment retains four canonical PRMs that corresponds to endophilin binding sites[36]. This region is highlighted as the boxed inset in (Figure 1A). Endophilin, hereafter referred to as EDP A1, comprises an N-terminal amphipathic helix (H0) and a BAR domain that generate and senses membrane curvature followed by a C-terminal SH3 domain that mediates binding to proline-rich partners (Figure 1A)[15, 38, 39]. For the GUV tubulation assay, fluorescently labelled EDPA1 or LPD 850-1250 was incubated with GUVs of lipid composition (DOPC: DOPE: DOPS: DHPE-MB 24.5:30:45:0.5 mole%). GUVs were considered as tubulated when they displayed thin, elongated cylindrical protrusions (≥1 µm) extending from their surface (Figure 1C). From the assay, GUVs were observed to exhibit different morphological categories such as, outer tubules, inner + outer tubules and inner tubules, and some GUVs without tubules, as shown in the schematic (Figure 1B). Both proteins exhibit membrane binding, as evident from their fluorescence enrichment along the vesicle surface, and they also exhibited tubulation (Figure 1C).

**Figure 1:**
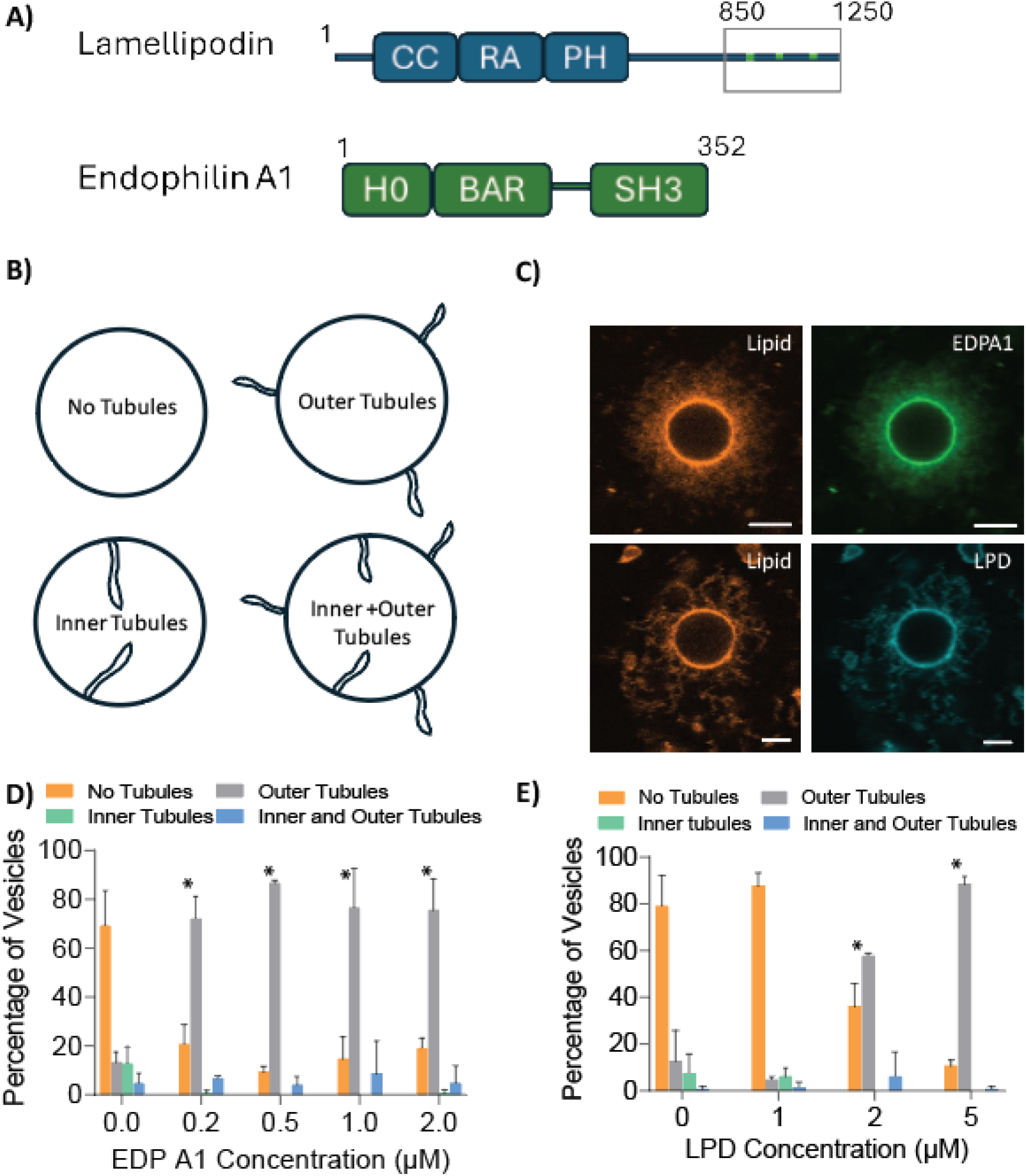
Endophilin and LPD drive curvature generation in GUV membranes. A) Schematic of functional domains of lamellipodin and endophilin A1. B) Schematic representation of GUV tubulation assay illustrating distinct vesicle morphologies: no tubules, outer tubules, inner tubules, inner + outer tubules. C) GUVs of lipid composition (DOPC: DOPE: DOPS; 24.5:30:45 mol%) labelled with fluorescent lipid (DHPE-MB, 0.5%), incubated with either endophilin A1 (0.5 μM EDP A1 with 20% AF-488 labelled, top) or LPD (5μM with 2% AF-647 labelled, bottom). D-E) Percentage of GUVs showing different curvature morphologies, including tubules, no tubules, inner tubules, and inner and outer tubules in the presence and absence of proteins. The data represents SD from three independent experiments. The number of GUVs analyzed in each set was 20-25. * *p* < 0.05 versus 0 µM; two-sided Dunnett-type adjusted Wald tests.

EDPA1 generated outward tubulation, with over 70% of GUVs exhibiting outer tubules at ≥ 0.2 µM (Figure 1D). LPD 850-1250 displayed concentration-dependent tubulation activity with approximately 57 and 88% of GUVs forming tubules at 2 and 5 µM, respectively, while most vesicles remained non-tubulated at 1µM (Figure 1E). Notably, in both cases, outer tubulation was the dominant phenotype. The occurrence of inner tubules or combined inner + outer tubules remained below 25% for both proteins across all tested concentrations indicating that inward curvature events were comparatively less prominent. In control experiments lacking either EDP A1 or LPD 850-1250, most of the GUVs remained spherical, with approximately 70 and 80% of vesicles, respectively, exhibiting no tubules. Collectively, these findings confirm the previously reported curvature generation capacity of endophilin[15, 29, 40] and further demonstrate that LPD is also capable of inducing membrane curvature at high concentration.

### Regulation of endophilin-mediated membrane curvature by SH3-PRM interactions with lamellipodin

Building on the understanding of how the individual proteins interact with membranes (Figure 1), we next asked how the cooperative interactions of EDPA1 and LPD 850-1250 influence membrane remodeling. To this end, GUVs were incubated with varying concentrations of EDPA1 and LPD 850-1250, while one protein was maintained at a constant concentration in each experiment and the concentration of the other one was varied. The resulting membrane morphologies were analyzed using confocal microscopy (Figure 2 A, C).

**Figure 2:**
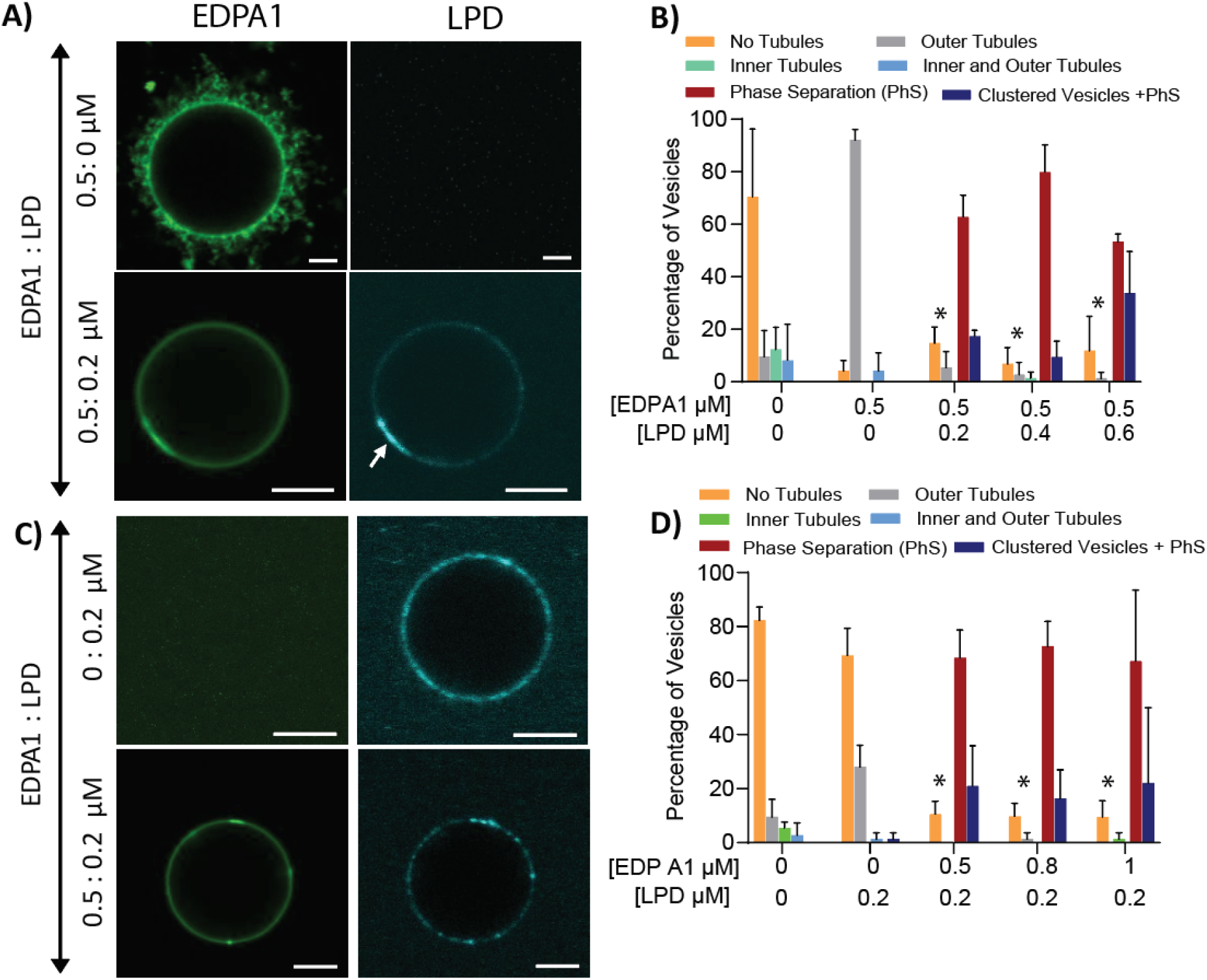
SH3-PRM interaction between endophilin and LPD regulates membrane remodeling and suppresses curvature generation. (A, C) Confocal images of GUVs (DOPC: DOPS: DOPE: MB-DHPE; 24.5:45:30:0.5 mole percent) with EDP A1 (green) and LPD (Cyan) at varying molar ratios. The GUVs were incubated with EDPA1 (0.5 μM) and either 0 or 0.2 μM of LPD (A) or with LPD (0.2 µM) and either 0 or 0.5 µM of EDP A1 (C). The scale bar represents 5 μm. (B and D) The percentage of GUVs exhibiting distinct morphologies-no tubules, outer tubules, inner and outer tubules, phase separation, and combination of phase separated clustered vesicles with tubules as a function of increasing LPD (0 to 0.6 μM) or EDPA1 (0 to 1 μM) concentrations at a constant EDPA1 (0.5 μM) or LPD (0.2 μM), respectively. The data represents SD from three independent experiments where 20-25 vesicles were examined per condition. * *p* < 0.05 versus 0 µM LPD (B) or 0 µM EPD (D); two-sided Dunnett-type adjusted Wald tests.

At a constant EDPA1 concentration (0.5 µM), increasing the amount of LPD 850-1250 from (0.2 - 0.6 µM) produced a marked change in GUV morphologies driven by SH3-PRM interactions between two proteins (Figure 2A). In the absence of LPD 850-1250, EDPA1 alone coats the GUV membrane relatively uniformly and predominantly generates outward tubulation (Figure 2A, Top row). However, the introduction of LPD 850-1250 modified this behavior. At LPD 850-1250 concentrations ranging from (0.2 µM -0.6 µM), the EDP-induced tubulation was suppressed, and GUVs displayed regions of protein-enrichment in both EDP A1 and LPD channels (Figure 2A, Second row). We hypothesize these domains to be consistent with protein-protein phase separation (see further below). Quantitative analysis shows that while >70% of vesicles exhibited outer tubules with EDPA1 alone, addition of 0.2 µM - 0.6 µM LPD reduced tubulation and increased the proportion of vesicles exhibiting protein phase separation to ≥50% (Figure 2B). These arise from the multivalent SH3-PRM network formed between EDPA1 and the highly PRM-enriched C-terminal region of LPD which facilitates protein clustering on the membrane[25, 41]. In addition, we examined the reverse conditions by keeping LPD at a constant concentration (0.2 µM) and titrating EDP (0.5 - 1µM) (Figure 2C,2D). Consistent with our observation in Figure 1E, the low concentration of LPD 0.2 µM alone did not induce membrane tubulation, and GUVs showed homogenous binding of LPD (Figure 2C, Top row). As EDPA1 concentration increased, the fraction of GUVs displaying outward tubules decreased sharply, while the proportion exhibiting phase-separated protein domains increased (Figure 2D). These results indicate that although EDPA1 alone is a strong driver for outward tubulation[15, 42], its curvature generation capacity is attenuated when LPD is present and able to bind through its PRM-rich C-terminal region.

Together, these complementary titration experiments demonstrate that the balance between membrane tubulation and phase separation of proteins is governed not merely by the individual membrane interaction activities of EDPA1 and LPD, but by their SH3-PRM dependent protein-protein interactions[36]. The resulting trends, suppression of tubules and emergence of protein-enriched domains, highlight a cooperative mechanism in which LPD modulates EDPA1-mediated membrane curvature through multivalent interactions[25].

### Multivalent interaction with lamellipodin shows conserved LLPS across endophilin A isoforms

Our group has recently shown that interactions between EDPA1 and LPD 850-1250 lead to the formation of phase-separated condensates in solution and protein clusters on the membrane [25, 41]. Endophilin’s LLPS is mainly driven by self-association of its N-BAR domain, whereas the SH3 domain mediates sequestration of PRM-containing binding proteins. Given that vertebrates express three endophilin isoforms A1, A2 and A3[30, 43, 44], we next sought to determine whether condensate formation is isoform-specific or conserved across the endophilin A isoforms. Although all three isoforms share an N-terminal H0 helix, a BAR domain and a C-terminal SH3 domain, the linker region connecting the BAR and SH3 domains differs in their length and composition (Figure 3A,B) suggesting that these variations may influence how each isoform engages with LPD 850-1250.

**Figure 3:**
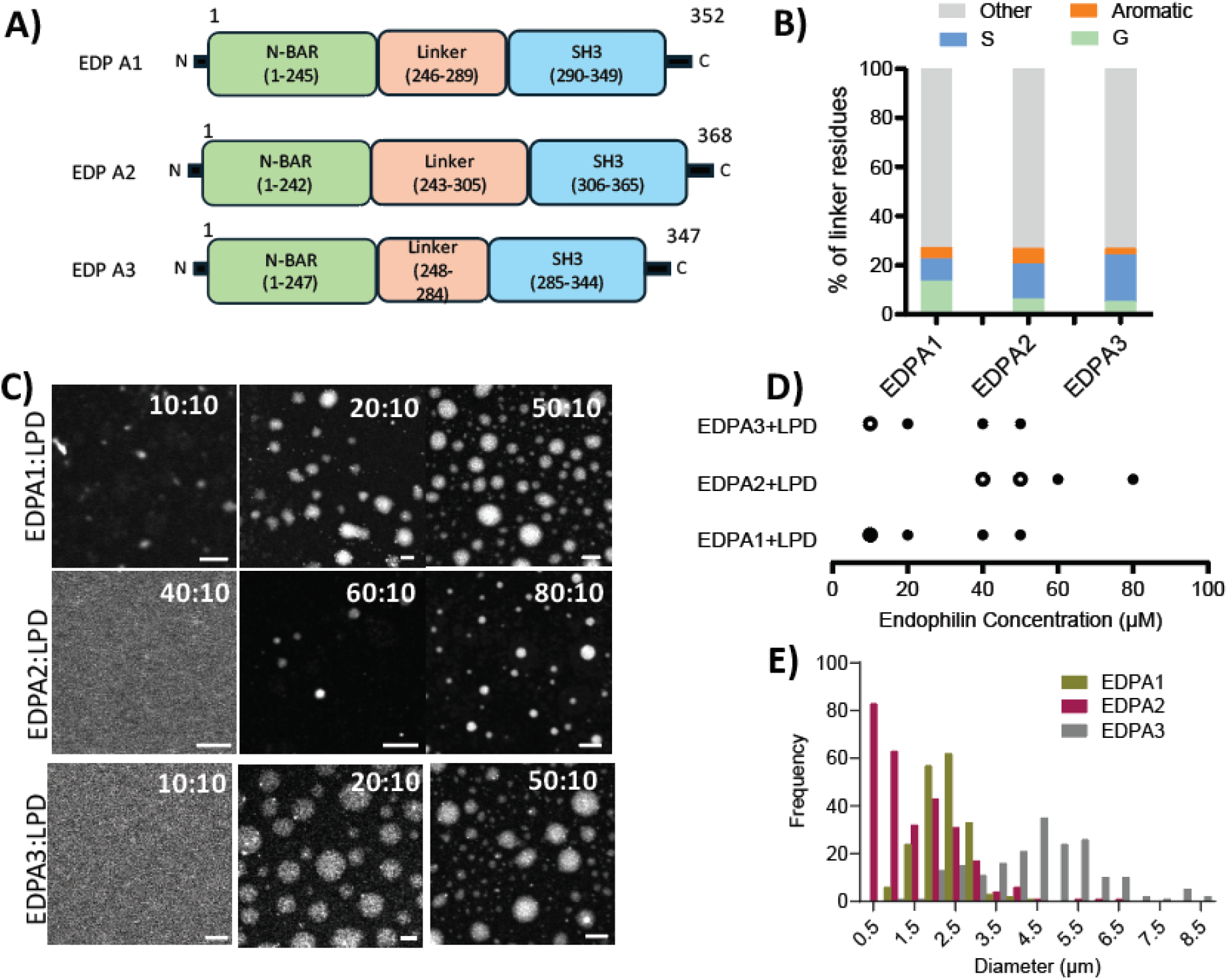
Isoform-dependent phase separation of endophilin A1, A2 and A3 with LPD. A) Schematic of endophilin A1, A2 and A3 isoforms showing conserved BAR domain, SH3 and variable linker region. B) Percentages of Serine (S), glycine (G), aromatic and other residues in the linker regions of EDP A1, A2, and A3 isoforms. C) Representative confocal images showing condensate formation of EDPA1, EDPA2 and EDPA3 with LPD at different molar ratios. Each row corresponds to the indicated EDPA1/A2/A3: LPD concentrations. The scale bar represents 5 μm. D) The phase diagram showing the condensate formation of EDP A1, EDPA2, and EDPA3 with LPD. The filled and unfilled circles indicate the presence and absence, respectively, of condensate formation while half-filled circles indicate conditions under which both droplets and puncta were observed. For panels C) and D), LPD was maintained at a constant concentration of 10 μM with 1% of Alexa Fluor 647 LPD and the number of independent experiments was *n* = 3. E) The size distribution curves of condensates formed by EDPA1: LPD, EDPA2: LPD and EDPA3: LPD.

Analysis of linker amino acid composition reveals the following differences among the three endophilin A isoforms (Figure 3B). The A1 linker contains the highest proportion of glycine and the lowest proportion of serine, whereas A3 shows the opposite pattern, with A2 exhibiting intermediate proportions of both residues. Aromatic residues constitute a small fraction of each linker, and residues outside these three categories account for the majority of the sequence in all isoforms. Thus, the linkers differ both in length and in the relative representation of glycine, serine, and aromatic residues, motivating a comparison of condensate formation across the endophilin A family.

To assess the phase separation capacity of the isoform with LPD, we examined condensate formation across a range of endophilin concentrations using confocal microscopy, while keeping LPD constant (Figure 3C). At the lowest equimolar condition (10:10 µM), no clear condensate formation was observed for EDPA2 or EDPA3, whereas EDPA1 exhibited a small number of droplets-like structures together with smaller punctate structures. Increasing the concentration to 20:10 µM led to the appearance of condensates for EDPA1 and EDPA3, whereas EDPA2 still showed no detectable condensates. At higher concentrations (50:10 µM for EDPA1 and EDPA3; 80:10 µM for EDPA2), all three isoforms formed condensates, indicating that multivalent interaction with LPD is sufficient to drive LLPS with modest isoform-dependent differences in the concentration thresholds. The phase diagram (Figure 3D) indicates the threshold concentration in the order EDPA1<EDPA3<EDPA2. In addition, we also quantified the size distribution of condensates formed by each isoform with LPD 850-1250 (Figure 3E).

### Peptide-mediated perturbation of SH3-PRM interaction leads to condensate dissolution

Endophilin interacts with several PRM-rich partners during endocytic trafficking, and Synaptojanin 1 is one of them[45, 46] The interaction depends on the SH3-domain binding to the COOH-terminal of the proline-rich domain of Synaptojanin 1, a region enriched in PRMs. Previous studies have shown that the PP19 peptide, which contains two PPXRP motifs from Synaptojanin 1, binds selectively to endophilin SH3 domains and blocks SH3-mediated interactions [47, 48]. Our earlier results demonstrate that condensate formation between endophilin A isoforms and LPD is driven by multivalent SH3-PRM interaction. Because PP19 retains the Synaptojanin’s essential SH3-binding sites, it can serve as a minimal model to probe how Synaptojanin-like PRM binding can modulate the stability of endophilin-LPD condensates. Using PP19 as a model peptide, we assessed its concentration-dependent effects on the stability of bulk phase condensates across all endophilin isoforms A1, A2 and A3. Our results show that PP19 destabilizes endophilin -LPD condensates in a concentration-dependent manner for all three endophilin isoforms (Figure 4A, B). In the absence of PP19, EDPA1 formed stable condensates with LPD consistent with our previous observations[25]. We further found that A2 and A3 isoforms also formed condensates with LPD under the same conditions. (Figure 4A, top row). At low concentration (0.5 µM) of PP19, the condensate number decreased noticeably for EDPA1 and EDP A2, although the condensates were still observed for all isoforms. At 5 µM of PP19, EDPA3-LPD condensates were largely dissolved, whereas EDPA1-LPD and EDPA2-LPD condensates persisted with lesser in number. At the highest PP19 concentration (25 µM), condensates from all isoforms dissolved resulting in a homogenous solution with no detectable condensates (Figure 4A). The partial phase diagram shows the presence and dissolution of EDPA1-LPD, A2-LPD and A3-LPD as PP19 concentration increases (Figure 4B). This trend aligns with the phase separation threshold defined earlier (Figure 3C), suggesting that isoform differences influence the formation and stability of the SH3-PRM network.

**Figure 4:**
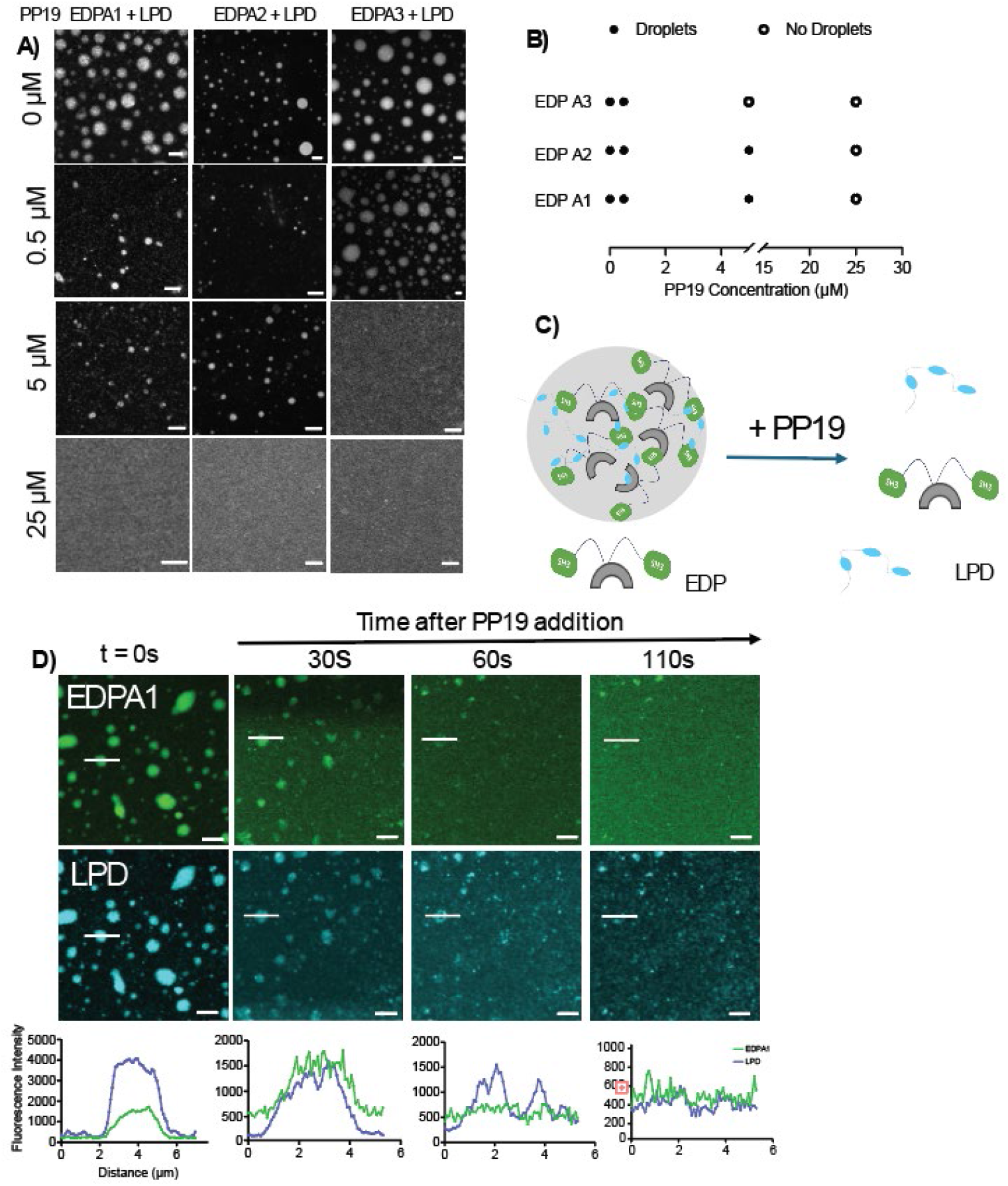
PP1G-mediated disassembly of protein condensates formed through SH3-PRM interaction. A) Representative confocal fluorescence images showing the PP19 mediated concentration-dependent disassembly of condensates formed by endophilin isoforms with LPD. LPD was maintained at 10 μM, with 1% AF-647 labelled LPD. The scale bar represents 5 μm. B) Phase diagram showing the concentration-dependent disassembly of condensates formed by EDP A1-LPD, EDP A2-LPD, and EDP A3-LPD condensates by PP19. Condensate presence is shown by filled circles and dissolution by open circles. Data were obtained from three independent experiments, *n* = 3. For both panels A and B, the protein concentrations were EDP A1 (50 µM), EDP A2 (80 µM) A3( 50 µM) and LPD (10 µM, 1% AF-647 labeled). C) Schematic representation of the proposed mechanism by which PP19 competes with the SH3-PRM interaction between endophilin and LPD, thereby disrupting the multivalent interaction leading to the condensate disassembly. D) Time-lapse confocal images show the kinetics of PP19-induced disassembly of EDPA1: LPD condensate. Images were acquired immediately before and after the addition of PP19 (100 µM). EDPA1 and LPD concentrations are 50 µM and 10 µM respectively. Progressive loss of fluorescence intensity indicates the dissolution of condensates. The scale bar indicates 5 μm. Fluorescence intensity line profile across the condensates (indicated by white lines) shows spatial colocalization of EDPA1 and LPD at *t* = 0 s and the gradual reduction of fluorescence intensity at later time points after PP19 addition.

To probe the kinetics of condensate disassembly, we monitored the time-resolved response of EDPA1-LPD condensates after PP19 addition. Immediately after PP19 exposure, condensates exhibit progressive shrinking and loss of contrast followed by complete dissolution within approximately 110 s (Figure 4C). The line intensity fluorescence profile across a representative droplet shows a rapid decrease in signal intensity for both EDPA1 and LPD (Figure 4D). Collectively, these results suggest that PP19 is capable of destabilizing EDP-LPD condensates by competing for SH3-domain binding, disrupting the multivalent SH3-PRM interaction network that stabilizes phase-separated assemblies (Figure 4E).

### PP1G rescues endophilin-driven membrane curvature by disrupting LPD-mediated SH3-PRM interaction

Building on our observation that PP19 disrupts EDP-LPD condensates, we next examined how this perturbation influences membrane curvature generation, extending our analysis from solution condensates to membrane remodeling. To that end, we incubated GUVs with EDPA1 and/or LPD, in the presence or absence of PP19, and imaged them using confocal microscopy. Figure 5A shows that EDPA1 generates outward membrane tubulation similarly on GUVs both in the absence and presence of PP19, indicating that PP19 binding to the SH3 domain does not seem to significantly modulate N-BAR-mediated curvature generation under our experimental conditions. When EDPA1 and LPD are co-incubated with GUVs, in the absence of PP19, EDP A1 and LPD form protein-enriched domains on the GUV surface without tubules, indicating that outward curvature generation by endophilin is suppressed when its SH3 domain is engaged by multivalent LPD PRM. However, when PP19 is added to the same co-incubated EDPA1 and LPD system, membrane tubulation reappears, confirming that PP19 binding to the EDPA1 restores curvature by disrupting multivalent SH3-PRM engagement interaction (Figure 5B). To determine whether PP19 modulates membrane curvature under our experimental conditions, we first examined its effect on GUVs in the absence of EDPA1 and LPD (Figure 5C). Incubation of GUVs with PP19 alone did not produce detectable membrane curvature and the majority of vesicles remained spherical. Consistent with its intrinsic curvature generation activity, EDPA1 retained its ability to generate outward tubulation of GUVs in the presence of PP19. Vesicles incubated with PP19 and LPD, largely retained their spherical morphology. The full distribution of vesicle phenotypes is shown in Figure S1. Together, these observations indicate that PP19 does not independently generate membrane curvature (Figure 5C).

**Figure 5:**
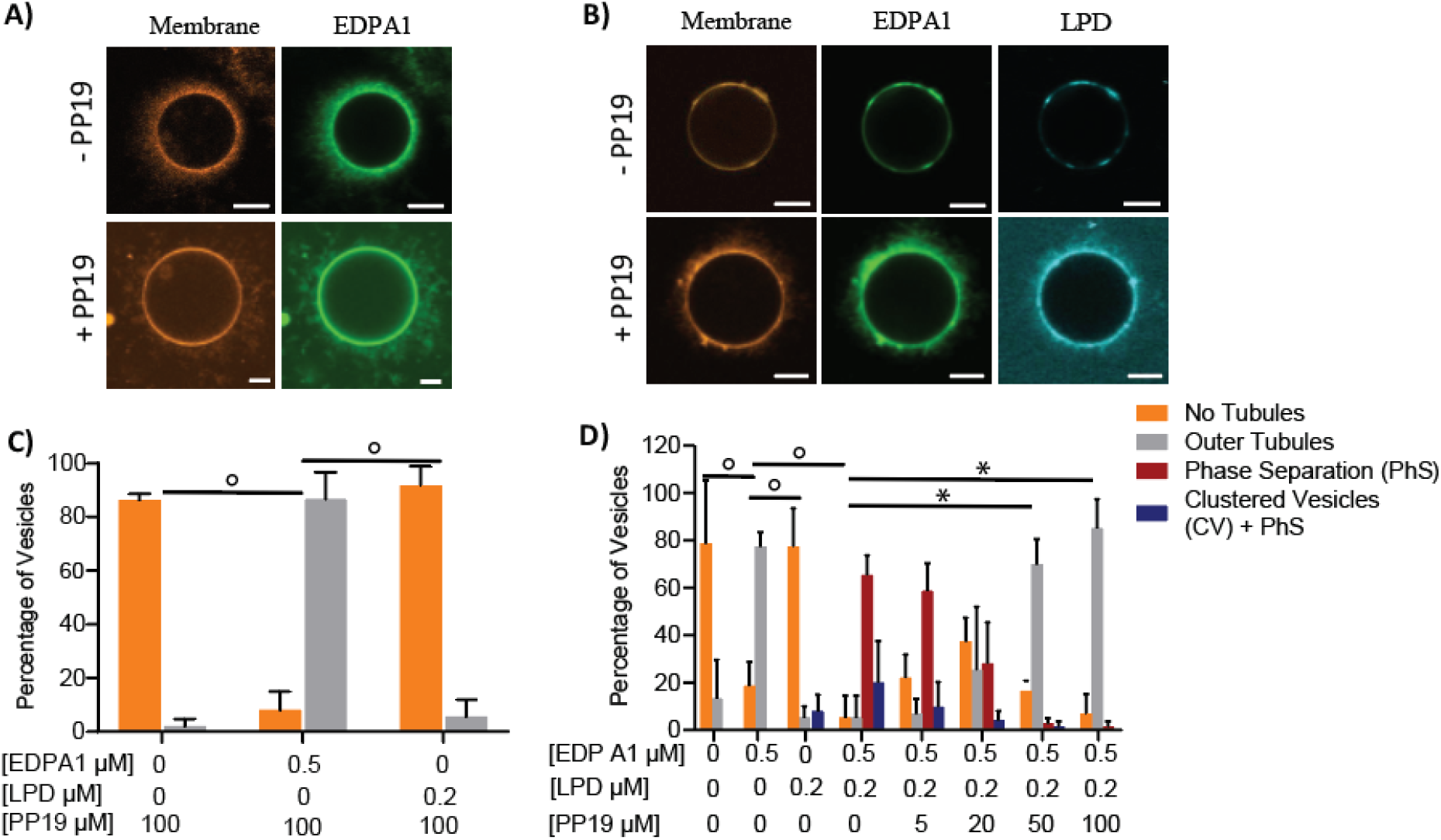
PP1G restores curvature generation by endophilin in the presence of LPD via SH3-PRM disruption. A) Representative confocal GUV images (DOPC: DOPS: DOPE: MB-DHPE; 24.5:45:30:0.5 mole percent) probing the curvature generation by EDPA1 (green) in the absence (top) and presence (bottom) of PP19 peptide (100 µM). B) GUVs incubated with EDP A1 (green) and LPD (Cyan) in the presence or absence of PP19 show curvature suppression (top panel, -PP19) and rescue the curvature respectively (bottom panel, +PP19, 100 µM). For both panels (A and B), the EDPA1 concentration was 0.5 µM. In panel B, the LPD concentration was 0.2 µM. GUV membranes were fluorescently labeled with DHPE-MB (0.5 mol%). The scale bar represents 5 µm. C) quantification of GUV tubulation in the presence of PP19 (100 µM) alone or in combination with EDP A1 (0.5 µM) or LPD (0.2 µM). D) Quantitative analysis of GUV morphologies under different conditions, such as in the absence and presence of only EDP A1, LPD, and EDPA1+LPD+PP19 (5 - 100 μM). Data represent mean ± SD from three independent experiments. The number of GUVs analyzed in each set was 20-25. ͦ *p* < 0.05; two-sided Wald test for the indicated comparison, * *p* < 0.05 versus 0 µM PP19; two-sided Dunnett-type adjusted Wald tests.

Figure 5D quantifies the concentration dependent effect of PP19 on membrane tubulation in GUVs co-incubated with EDP and LPD. To establish baseline membrane remodeling behaviors, we first analyzed GUV morphology in single and two-component systems prior to PP19 titrations. These controls allowed us to determine how each protein individually and in combination with other components influences membrane curvature.

For the single-component systems, GUVs were incubated separately with EDP isoforms, LPD, and PP19. In the absence of proteins, GUVs exhibited minimal tubulation (13% outer tubules, 78% non-tubulated vesicles). To minimize background curvature, we selected GUV batches exhibiting a low outer-tubule fraction (<25%) for all subsequent single, two-component, and PP19 concentration-titration experiments. EDPA1 on GUVs consistently generated outward membrane tubules (77%) and minimal non-tubulated vesicles (18%). LPD only and PP19 only controls exhibited non-tubulated vesicle fractions of (76% and 86% respectively), with the tubulated fraction of 5% and 2%. These controls confirm that curvature generation is specific to EDP alone, whereas LPD or PP19 independently do not generate curvature (Figure 5C and D).

We next examined two (protein-) component systems, to determine how interactions between these components influence membrane remodeling. The two-component system of EDPA1 and LPD consistently showed tube suppression, with most vesicles exhibiting protein phase separation of 65% and tubulated fractions of 5% (Figure 5D). In the EDPA1+ PP19 system, the vesicles are predominantly tubulated (85%), confirming that PP19 addition did not alter EDPA1’s intrinsic curvature generation property under our experimental conditions. Also, the LPD+PP19 system showed the vesicles were predominantly non-tubulated (88%) indicating that PP19 presence neither activates nor amplifies the curvature generation (Figure 5C).

Next, we introduced increasing concentrations of PP19 (5-100 µM) into GUV dispersions, co-incubated with EDP-LPD, to test the concentration-dependent reversal of tube suppression. At 5 µM of PP19, in comparison to EDP-LPD binary component system (5% outer tubules, 65.33% phase separation), there is no considerable increase in outward tubule recovery (6.8 % outer tubules, 58.78% phase separation, 22% no tubules). While at 20 µM, PP19 enhanced tubulation (25% outer tubules, 28% phase separation, 37% no tubules). At 50 µM, the system shifted predominantly to outer tubules (69% outer tubule GUVs) with only 2.7% and 16.4% protein phase separation and no tubule categories remaining and at 100 µM PP19, phase separated vesicles were close to absent, contributing 1% of the total vesicle population. Instead, under these conditions, outward tubules dominated (85% outer tube positive vesicles), and the ‘no tubule’ population declined to 6%. The last two concentrations show statistically significance release of tubulation suppression (via logistic regression with Dunnett adjustment for multiple testing, see Methods section). All other GUV categories such as inner tubules, inner and outer tubules, phase separation + clustered vesicles and tubes remained consistently low across the PP19 concentration and did not exceed 25% of the population (Figure S2). These data establish a progressive transition from tubule suppressed state in EDP-LPD state to outward curvature recovery with increasing PP19 peptide levels.

The reversible switch between tube suppressed, phase separated state and the curvature restoration state supports a model (see Figure 9) where SH3-dependent recruitment of multivalent LPD drives membrane remodeling away from N-BAR mediated curvature, while PP19 restores curvature by displacing PRM-dependent suppression within the multivalent network.

### PP1G reverses LPD-mediated suppression of Endophilin A2/A3 driven membrane tubulation

We next examined whether PP19 could reverse LPD-mediated suppression of membrane tubulation by EDPA2 and EDPA3. In these experiments, EDP and LPD were first incubated with GUVs followed by the addition of PP19. This order of addition allowed us to test whether PP19 could relieve an already established LPD-dependent suppression of endophilin-mediated membrane modelling. Having first estimated the concentration threshold required for EDPA1 to generate outward curvature, we next asked whether this threshold differs across the other endophilin isoforms. Relative to the lower outer tubulation threshold observed for EDPA1 (≥0.5 µM), both EDPA2 and EDPA3 require higher concentrations to induce curvature. EDPA2 showed a threshold with curvature detectable around ∼1 µM and increasing substantially at 5 - 10 µM (Figure S2A). In contrast, EDPA3 exhibited the weakest curvature activity, showing an onset of outward tubules at 5 µM and pronounced tubulation at 10 - 25 µM (Figure S2B). Together, these trends indicate that the tubulation thresholds of the endophilin isoforms increase in the order of EDPA1 < EDPA2 < EDPA3 (Figure 1D, S2A and B).

Because the endophilin A isoforms exhibited different concentration thresholds for membrane curvature generation, each isoform was tested at a concentration sufficient to produce robust tubulation under its respective control condition. Under these isoform-specific baseline conditions, PP19 produced concentration-dependent changes in membrane remodeling across the endophilin isoforms. EDPA2 responded to low PP19 concentrations (20 µM) and restored outward curvature (>50%) from the tube-suppressed state (Figure 6A). In contrast, both EDPA3 and EDPA1 required a higher PP19 level (50 µM) to achieve a comparable recovery of curvature (Figure 6B and Figure 5D). The comparison of PP19 responses across EDPA isoforms is shown in Figure 6D. In addition, both EDPA2-LPD and EDPA3-LPD controls showed a tube suppressed state consistent with EDPA1-LPD system. Curvature suppression was calculated as relative to the basal endophilin induced curvature. Comparable suppression levels across EDP isoforms indicate that multivalent SH3-PRM interactions similarly inhibit intrinsic curvature generation across isoforms (Figure 6C). Moreover, PP19 did not alter the intrinsic curvature generation capacity of EDPA isoforms

**Figure 6:**
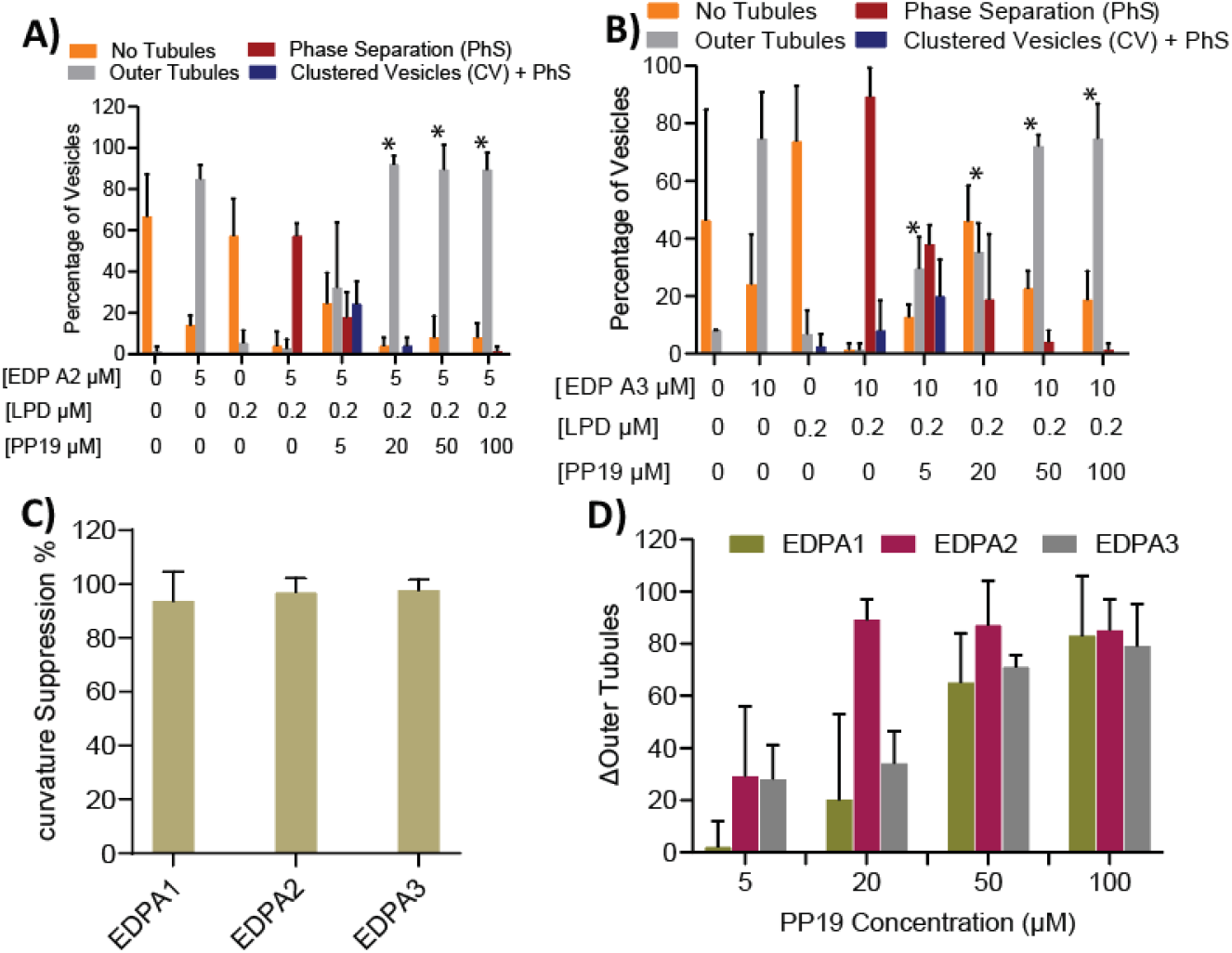
Comparable PP1G mediated rescue of membrane curvature across endophilin A2 and A3 isoforms. A, B) Quantitative plots of the concentration-dependent restoration of membrane curvature in GUVs (DOPC: DOPS: DOPE: MB-DHPE; 24.5:45:30:0.5 mole percent) by PP19 in the presence of LPD for EDPA2 and EDPA3, respectively. C) LPD mediated suppression of intrinsic endophilin-driven membrane curvature across isoforms. D) PP19 mediated recovery of LPD-suppressed membrane curvature across endophilin isoforms. For both panels C and D, the concentrations were EDP A1 (0.5 µM), EDPA2 (5 µM), EDPA3, (10 µM) and LPD (0.2 µM). * *p* < 0.05 versus 0 µM PP19; two-sided Dunnett-type adjusted Wald tests.

### PRM7 recapitulates multivalency-dependent condensate formation and curvature suppression, both restored by PP1G

To determine whether repeated SH3-binding motifs are sufficient to reproduce the effects of lamellipodin on condensate formation and membrane curvature, we examined a simplified multivalent construct, PRM7. This construct contains seven tandem repeats of a single endophilin-binding PRM from lamellipodin, separated by flexible linkers, allowing us to assess the contribution of PRM multivalency independently of the remaining lamellipodin sequence. Consistent with LPD, PRM7 formed condensates with EDPA1 in the absence of PP19, showing that tandem PRMs are sufficient to drive phase separation (Figure 7A). As PP19 was titrated into the system, condensates disappeared only at substantially higher concentrations than with LPD-EDPA1 (Figure 7A, 7B). The LPD-driven condensates were destabilized at 25 µM PP19, whereas PRM7 condensates remained intact across this range and disassembled at 100 µM, indicating that increased PRM valency amplifies the stability of the SH3-PRM interaction network.

**Figure 7:**
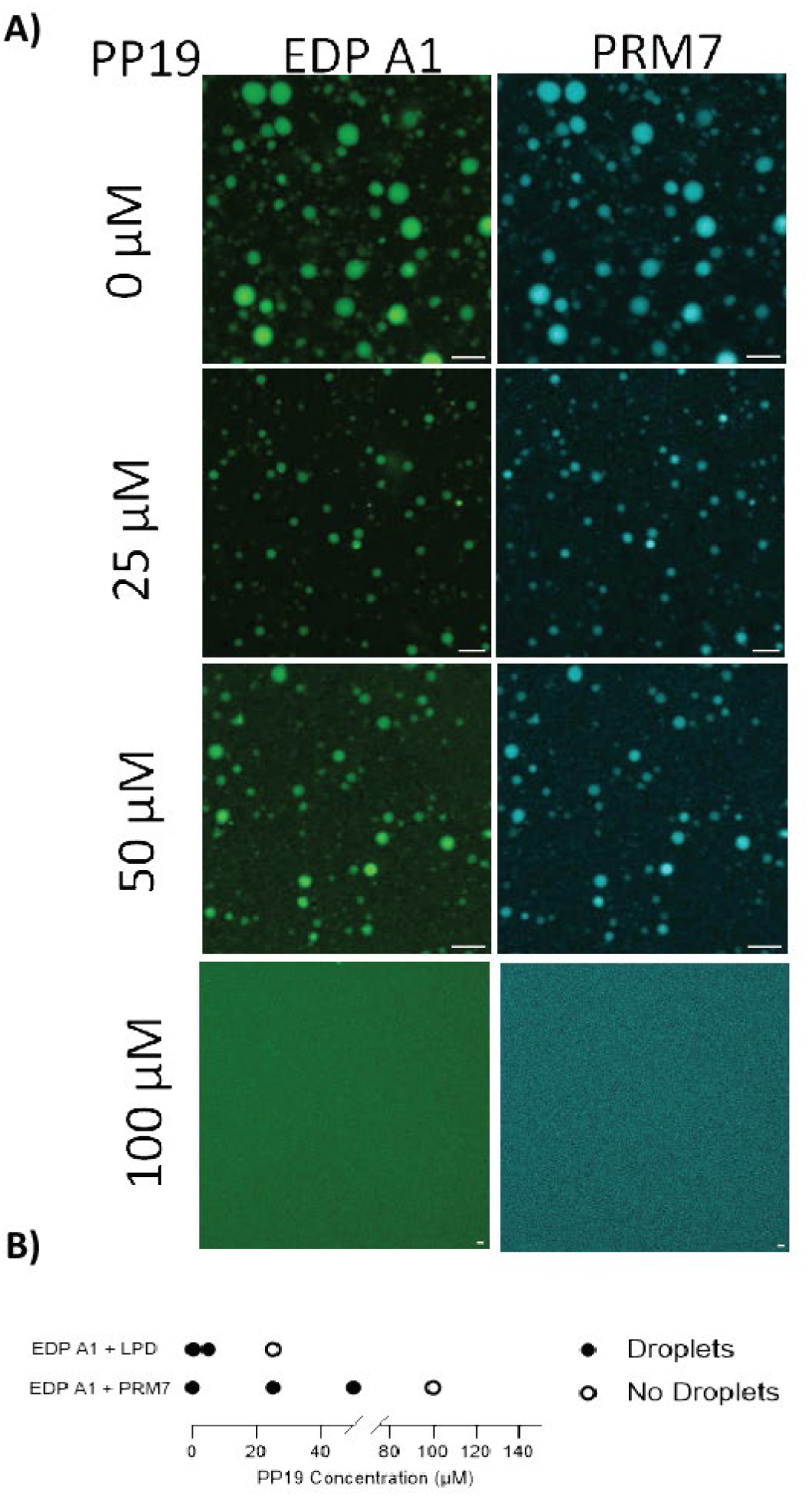
PP1G disrupts endophilin-PRM 7 condensates in a concentration-dependent manner. A) Phase separated condensates formed by EDPA1 (5 µM, 2% AF-488 labeled) and PRM7 (20 µM, 0.5% AF-647 labeled) undergo disassembly at higher PP19 concentration (100 μM). B) The phase boundary of EDPA1-PRM7 (5:20 µM) and EDPA1-LPD (50:10 µM) condensates in the presence of varying PP19 concentrations.

So far we have verified that multivalent PRMs are sufficient to generate condensates, with PRM7 forming more PP19 resistant condensates than LPD. To determine whether multivalency exerts a similar influence on membrane curvature, we next turned to LPD-C4, a monovalent PRM element containing the KPPPxPxR sequence of the PRM7 construct that retains a single SH3 binding motif but lacks multivalent interaction capacity (Figure 8C). This allowed us to directly test whether curvature suppression depends on multivalent SH3-PRM engagement. Consistent with PP19, our results show that the monovalent LPD C4 does not alter endophilin-mediated membrane curvature. The incubation of EDPA1 alone with GUVs generated outward tubules, whereas co-incubation with LPDC4 neither suppressed tubulation nor generated protein-enriched regions (Figure 8A). Quantitative analysis showed that the percentage of tubulated vesicles in EDPA1+LPDC4 was comparable to that in EDPA1 alone (Figure 8B). These findings show that a single SH3-binding motif is insufficient to interfere with N-BAR-driven curvature generation, and that multivalent PRM engagement is required for curvature suppression, as seen with LPD. We therefore next asked, whether PRM7 composed of seven LPDC4 repeats, could reconstitute curvature suppression. When incubated with GUVs, EDPA1 alone generated outer tubules, whereas co-incubation with PRM7 shifted the system to a curvature-suppression state (Figure 8D). The quantification confirmed substantial reduction in outer tubules and increase in protein-phase separated state in the presence of PRM7 (Figure 8E). Next, to test whether PRM7 mediated curvature suppression is reversible through the disruption of SH3-PRM interaction, we titrated PP19 into the EDP-PRM7 system. At low PP19 concentrations, protein enriched regions persisted, and outer tubules remains suppressed (Figure 8E) However, at higher PP19 concentrations, the outer tubules was restored (Figure 8D,8E).

**Figure 8:**
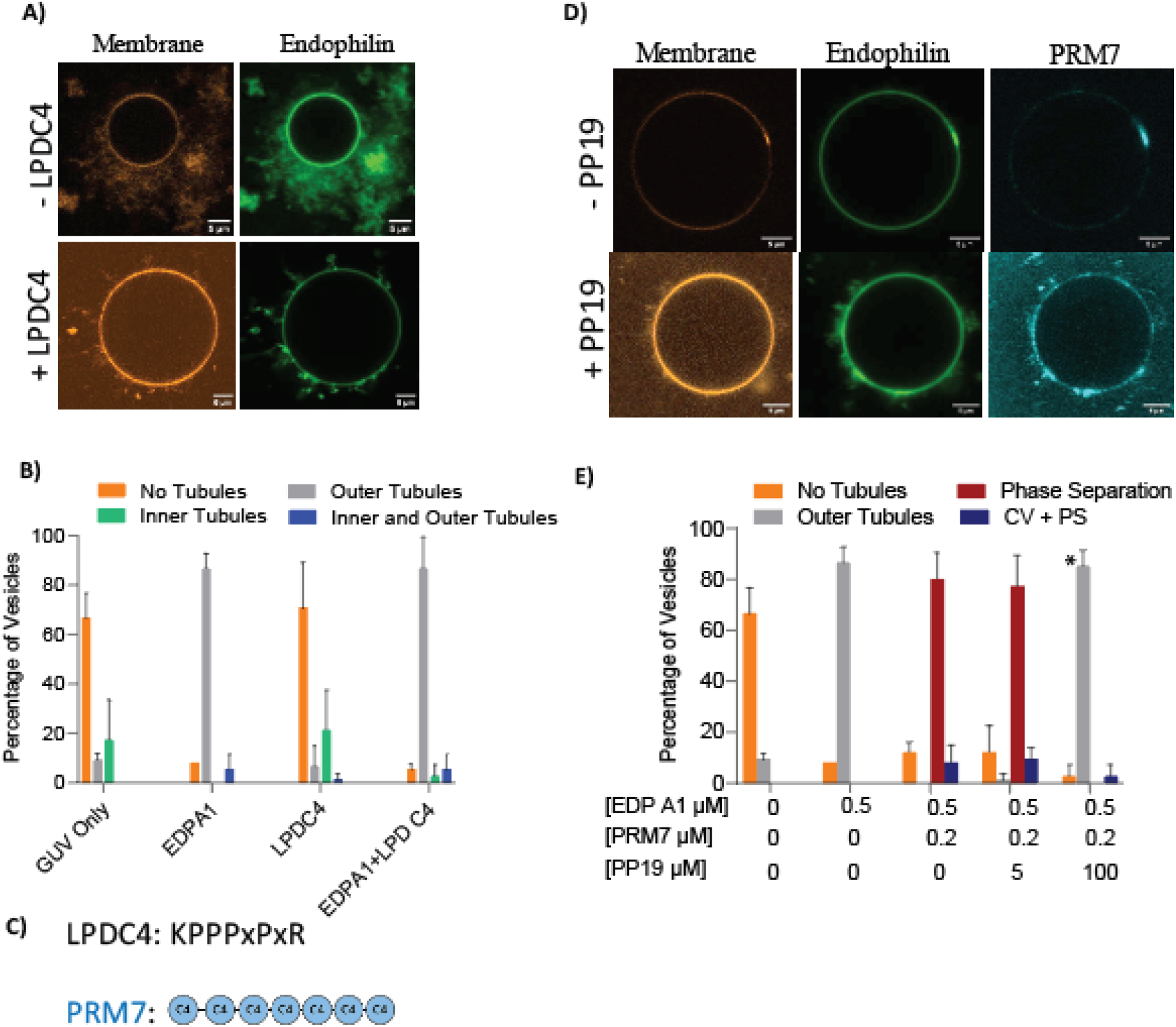
PRM7 reconstitution shows SH3–PRM-mediated curvature regulation by multivalency is rescued by PP1G. All GUVs had composition DOPC: DOPS: DOPE: MB-DHPE; 24.5:45:30:0.5 mole percent. A) Representative confocal images of GUVs incubated with EDPA1 (0.5 µM, 20% AF-488 labelled) in the absence (top) or presence (bottom) of LPD C4 (0.2 µM). B) Quantitative plot showing curvature generation with EDPA1 (0.5 µM) in the presence and absence of LPDC4 (0.2 µM). C) Schematic representation of the LPDC4 motif and the multivalent PRM7 containing seven LPDC4 repeats. D) Confocal fluorescence images of GUVs incubated with EDPA1 (0.5 µM, 20% AF-488 labelled) and PRM 7 (0.2 µM, 50% AF-647 labelled) in the absence (top) or presence (bottom) of PP19 100 µM showing curvature suppression and restoration respectively. For panels A and D, GUVs were labeled with the fluorescent lipid DHPE-MB (0.5 mol%). The scale bar represents 5 μm. E) Quantitative analysis of GUV morphologies showing curvature restoration in the presence of PP19 at a higher concentration of 100 µM. The data represent the mean ± SD from three independent experiments. The number of GUVs analyzed in each set was 20-25. * *p* < 0.05 versus 0 µM PP19; two-sided Dunnett-type adjusted Wald tests.

Taken together, our results demonstrate that multivalent PRM interactions are required for both condensate formation and inhibition of N-BAR-mediated curvature, with LPD and PRM7 providing the necessary valency and LPDC4 confirms that monovalent SH3 binding alone is insufficient to modulate endophilin’s membrane remodeling output.

## Discussion

Here we have shown that multivalent interactions between endophilin and lamellipodin, the core molecular components of fast endophilin mediated endocytosis (FEME) play a regulatory role in membrane remodeling. SH3-PRM driven multivalency promotes the formation of a protein rich phase on the membrane that constrains the intrinsic curvature generation capacity of endophilin. Our data show that curvature generation in FEME is therefore not dictated simply by the presence of N BAR proteins, but by the valency-dependent organization and phase behavior of their interactions.

Membrane remodeling during endocytosis requires the coordinated action of proteins that both sense and generate membrane curvature while simultaneously engaging a complex network of interaction partners. Endophilin occupies a central position in this process through its N-BAR domain, which drives membrane deformation, and its SH3 domain, which recruits numerous proline-rich binding partners. Although previous studies have established that endophilin and its interaction partners can undergo liquid-liquid phase separation and form membrane-associated protein assemblies, the functional relationship between condensate formation and membrane remodeling has remained unclear. In the present study, we demonstrate that multivalent SH3-PRM interactions between endophilin and lamellipodin establish a regulatory mechanism that suppresses membrane tubulation while promoting the formation of protein-rich condensed phases. Conversely, disruption of these interactions by the synaptojanin-derived peptide PP19 dissolves condensates and restores endophilin-mediated membrane curvature generation. Together, these findings identify multivalent protein interactions as a molecular switch that controls the balance between phase separation and membrane remodeling (Figure 9).

**Figure 9:**
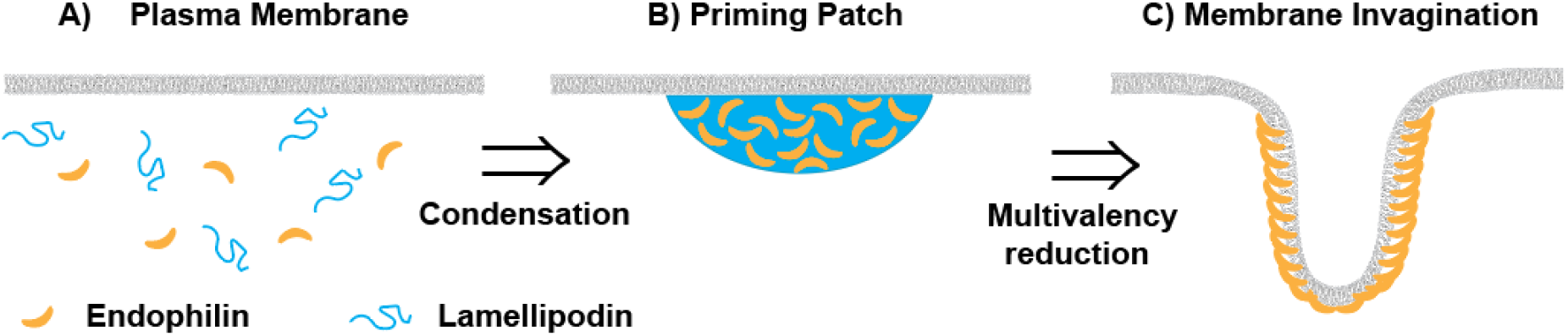
Model showing the regulatory role of PP1G in restoring endophilin curvature generation. (A) Unprimed plasma membrane: EDP and LPD are dispersed in the cytoplasm. (B) Multivalent EDP-LPD interactions form a membrane-associated priming patch that restrains curvature. (C) PP19 disrupts these interactions and endophilin-mediated membrane curvature generations is restored.

A central observation of this work is that endophilin-mediated membrane tubulation and protein condensation appear to represent competing organizational states. Endophilin alone efficiently generates membrane tubules on anionic GUV membranes, consistent with its well-established role as a membrane curvature generator. In contrast, when lamellipodin is present at concentrations sufficient to engage endophilin through multiple SH3-PRM interactions, tubulation is strongly suppressed and replaced by the appearance of membrane-associated protein-rich domains. This behavior was observed regardless of whether endophilin or lamellipodin was titrated into the system, indicating that the effect arises from the multivalent interaction network itself rather than from changes in the concentration of either individual component. These findings suggest that assembly of a dense SH3-PRM interaction network redirects endophilin from a curvature-generating state into a condensed state characterized by extensive protein-protein interactions.

The molecular mechanism by which condensate formation suppresses membrane curvature generation remains to be determined. One possibility is that formation of a dense multivalent network restricts the translational or rotational mobility of membrane-bound endophilin molecules, thereby reducing their ability to undergo the cooperative organization required for tubule formation. Alternatively, condensate formation may alter the local arrangement or effective concentration of N-BAR domains in a manner that favors protein-protein interactions over membrane deformation. A further possibility is that clustering of endophilin and lamellipodin generates membrane-associated phases with distinct physical properties that oppose the collective membrane remodeling activity of BAR-domain assemblies. Although our experiments do not yet distinguish among these possibilities, they clearly establish that engagement of endophilin in a multivalent SH3-PRM network is sufficient to suppress its intrinsic membrane-remodeling activity.

The observation that PP19 reverses both condensate formation and curvature suppression provides further support for this model. PP19 competes for binding to the endophilin SH3 domain and thereby disrupts the multivalent interaction network that stabilizes endophilin-lamellipodin assemblies. Importantly, PP19 did not alter the intrinsic ability of endophilin to generate membrane curvature when endophilin was examined in isolation. Instead, its principal effect was to restore tubulation in systems where curvature generation had previously been suppressed by multivalent interactions. We note that the concentrations for PP19 used here are higher than the cytosolic concentrations of typical endocytic accessory proteins. For those, as opposed to the PP19 model peptide, additional protein and lipid interaction domains may amplify binding interactions. Our findings demonstrate that the inhibitory effect on membrane remodeling is not a consequence of direct regulation of the N-BAR domain but rather emerges from SH3-dependent network formation. The rapid dissolution of condensates following PP19 addition further emphasizes the dynamic nature of these assemblies and suggests that transitions between condensed and membrane-remodeling states can occur on functionally relevant timescales. We emphasize that this mechanism of the orchestration of membrane shape transition through the tuning of multivalency is not limited to a mechanism based on competitive binding interactions. It is likely that multivalency tuning can be accomplished rapidly in signaling networks through reversible posttranslational modifications, including phosphorylation / de-phosphorylation.

An important finding of this study is that this regulatory behavior is conserved across the endophilin A family. Although endophilin A1, A2, and A3 exhibited modest differences in their thresholds for condensate formation, membrane tubulation, and PP19 sensitivity, all three isoforms displayed the same qualitative response: multivalent interaction with lamellipodin promoted condensate formation and suppressed membrane curvature generation, whereas disruption of these interactions restored tubulation. The conservation of this behavior despite differences in linker length and sequence composition suggests that the underlying mechanism arises from architectural features shared across the endophilin family. More broadly, these results imply that coupling between multivalent interaction networks and membrane remodeling may represent a general property of endophilin-dependent trafficking pathways rather than the behavior of a single isoform.

The relative glycine, serine, and aromatic residue contents of the linkers do not show a consistent relationship with the observed onset of phase separation (Figure 3B, D), arguing against a simple explanation based on these compositional features alone. In particular, A1 and A3 differ substantially in their glycine–serine balance but form condensates within the same tested concentration interval, whereas A2 requires higher concentrations. Linker length provides an alternative hypothesis: A2 possesses the longest linker, which could alter domain organization or increase the steric and solvation costs of assembly. This possibility is consistent with previous evidence suggesting that disordered linkers can restrain BAR-domain-driven condensation [25] and with theoretical work demonstrating that linker length and solvation properties modulate phase separation in multivalent proteins [49]. However, the isoforms also differ outside their linkers, and targeted linker modifications will be required to establish the contribution of linker length.

The experiments using PRM7 and LPD-C4 provide additional mechanistic insight into the role of multivalency. Whereas the monovalent LPD-C4 construct failed to suppress endophilin-mediated tubulation, the multivalent PRM7 protein recapitulated both condensate formation and curvature suppression. Moreover, PRM7 assemblies exhibited increased resistance to PP19-mediated dissolution compared with lamellipodin-derived assemblies. Together, these findings demonstrate that simple occupancy of the SH3 domain is insufficient to regulate membrane remodeling. Instead, network connectivity and interaction valency emerge as critical determinants of both condensate stability and functional regulation. This conclusion aligns with broader principles that have emerged from studies of biomolecular condensates, in which the number and arrangement of interaction motifs often govern both phase behavior and biological function.

Our findings also have implications for the regulation of fast endophilin-mediated endocytosis. FEME requires the coordinated recruitment of endophilin, lamellipodin, and numerous additional SH3-binding partners to discrete membrane sites. Previous work has suggested that clustering and phase separation may contribute to the spatial organization of these assemblies. The results presented here suggest a more nuanced view in which condensate formation not only concentrates proteins but also modulates their mechanical activity. In this framework (see Figure 9), multivalent interactions could stabilize a protein-rich pre-endocytic assembly state, whereas competition from additional binding partners or changes in local protein composition could trigger a transition toward active membrane remodeling. Such a mechanism would provide cells with a means to couple biochemical assembly of the endocytic machinery to the physical process of membrane deformation.

Several limitations should be considered when interpreting the present results. The experiments were performed using purified proteins and model membranes, which allow direct interrogation of molecular mechanisms but necessarily simplify the cellular environment. Additional endocytic proteins, membrane heterogeneity, cytoskeletal interactions, and active cellular processes may influence the balance between phase separation and membrane remodeling in vivo. Furthermore, while our results establish a strong relationship between condensate formation and suppression of membrane curvature generation, they do not yet reveal the structural basis of this coupling. Future studies combining quantitative biophysical measurements with high-resolution structural and dynamical approaches will be required to determine how multivalent protein assemblies alter the membrane-remodeling activity of BAR-domain proteins.

## Conclusion

We demonstrate that multivalent SH3-PRM interactions between endophilin and lamellipodin regulate membrane remodeling by shifting endophilin from a curvature-generating state into a phase-separated protein-rich state. This behavior is conserved across endophilin isoforms, depends critically on interaction valency, and can be reversibly controlled by competitive SH3-domain ligands. These findings establish a direct functional link between biomolecular condensation and membrane remodeling and suggest that phase separation can serve not only as a mechanism for molecular organization but also as a regulator of the mechanical activities that drive membrane trafficking.

## Acknowledgments

We acknowledge funding from NIH grant GM097552 and the UPenn Research Foundation, and helpful discussions with Samsuzzoha Mondal and Karthik Narayan.

## Supplementary Figures

**Figure S1:**
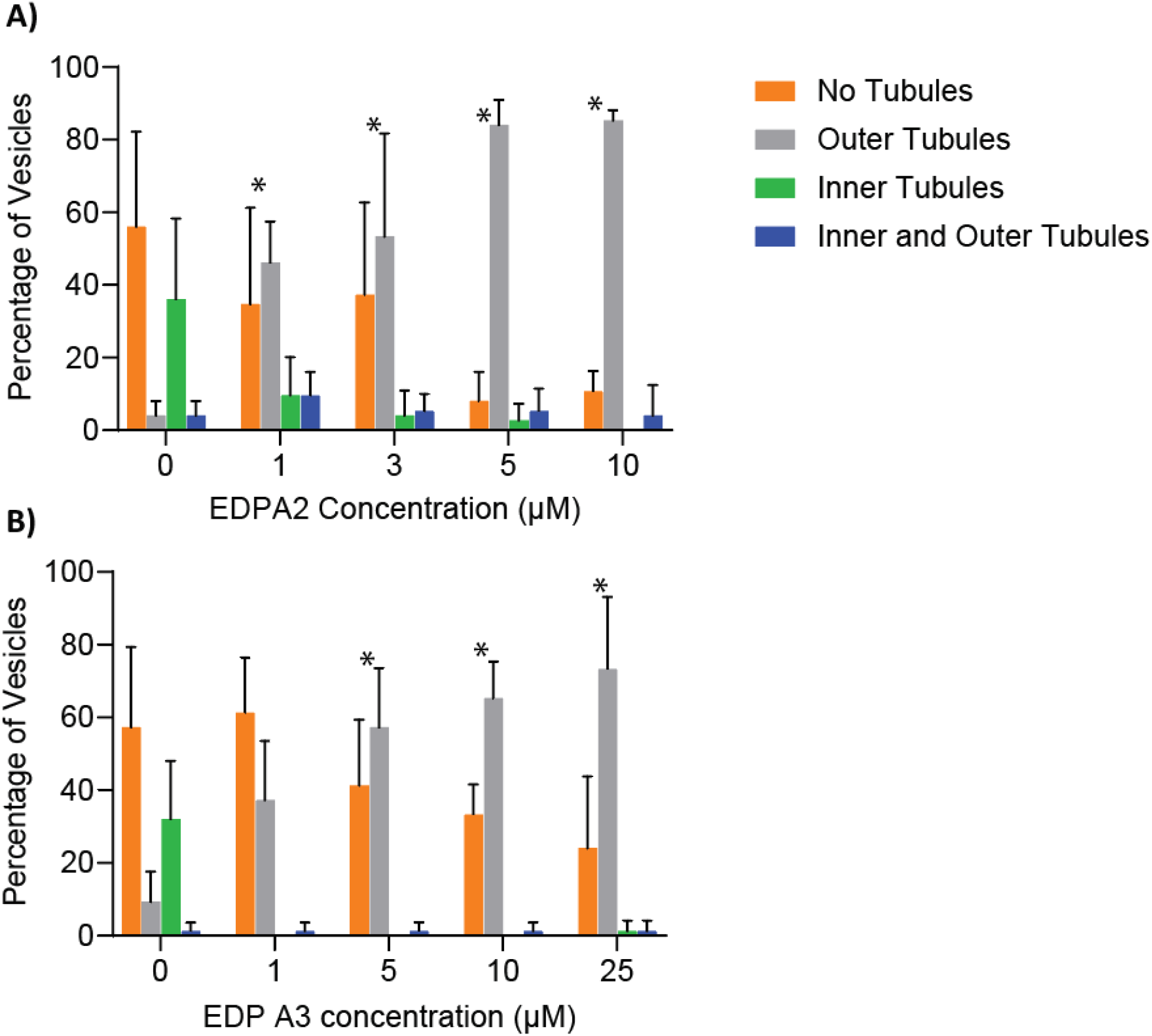
A, B) Quantitative analysis of curvature generation in GUVs (DOPC: DOPS: DOPE: MB-DHPE; 24.5:45:30:0.5 mole percent) by EDP A2 and EDPA3 at increasing protein concentrations. Data is presented as mean ±SD from three independent experiments (*n* = 3) with 20-25 GUVs analyzed per experiment. * *p* < 0.05 versus 0 µM EDP; two-sided Dunnett-type adjusted Wald tests.

**Figure S2:**
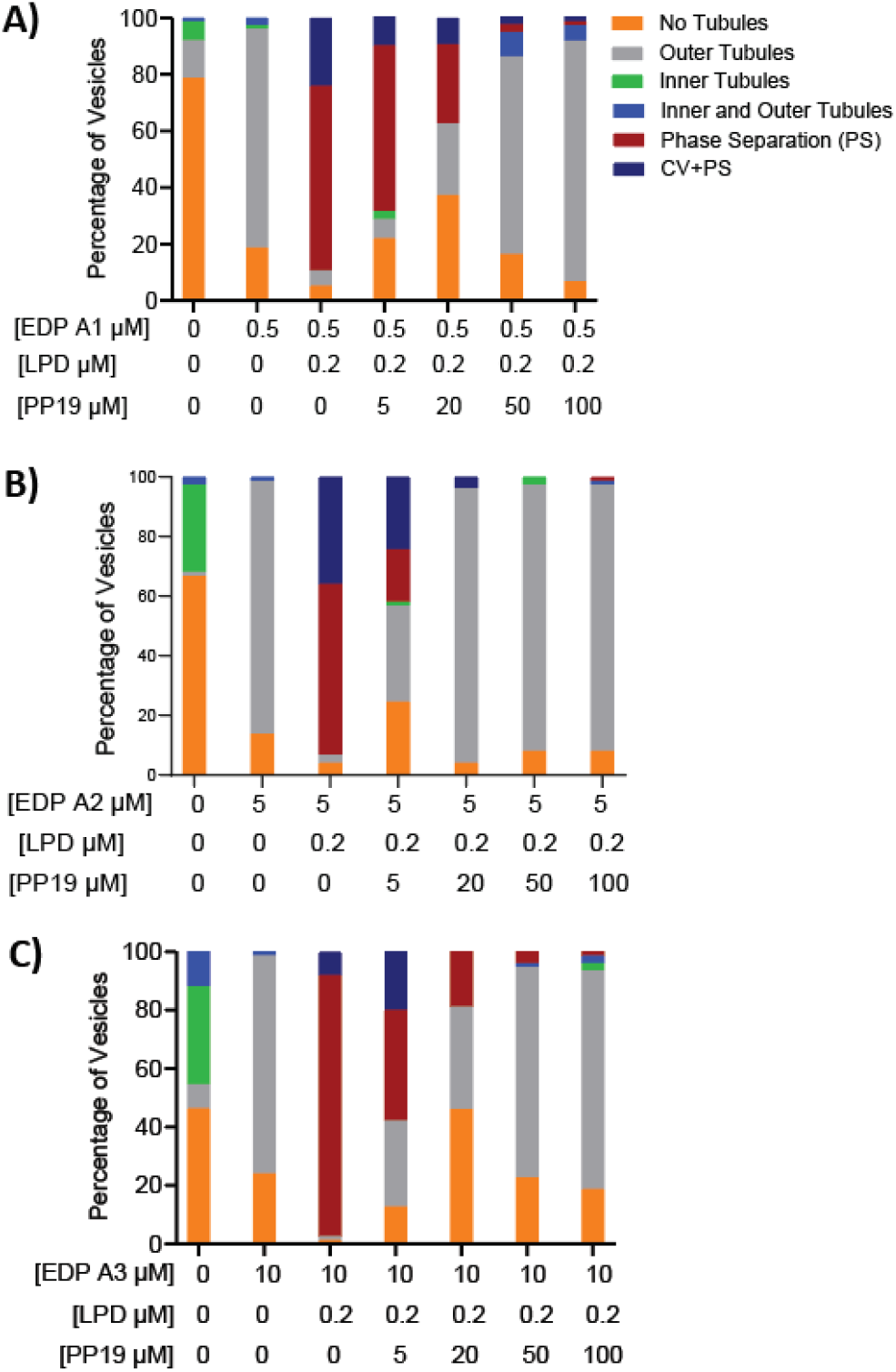
A, B, and C) Comprehensive vesicle (DOPC: DOPS: DOPE: MB-DHPE; 24.5:45:30:0.5 mole percent) phenotype distribution across the indicated EDP isoforms, LPD and PP19 conditions. Stacked bars represent the mean percentage from three independent experiments with 20-25 GUVs analyzed per experiment.

**Figure S3:**
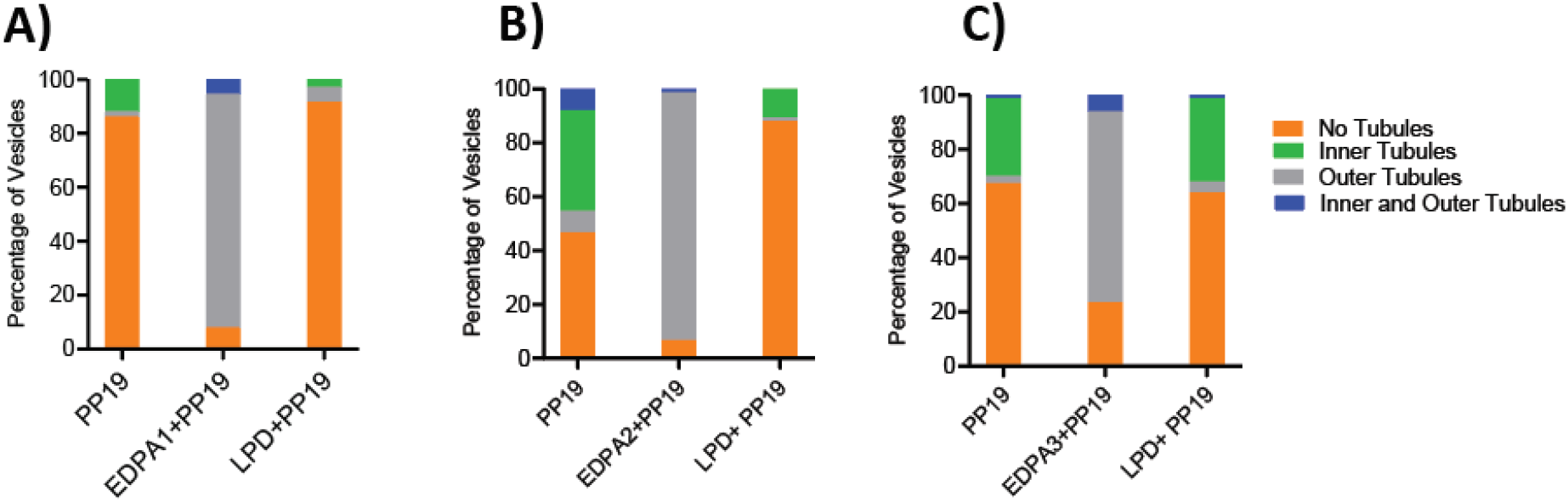
A, B and C) Complete vesicle morphology (DOPC: DOPS: DOPE: MB-DHPE; 24.5:45:30:0.5 mole percent) classification of PP19 mediated curvature modulation in the presence of endophilin isoforms alone or LPD. The protein concentrations were EDP A1 (0.5 µM), EDP A2 (5 µM), EDP A3 (10 µM), LPD (0.2 µM) and PP19 (100 µM). Stacked bars represent mean percentage from three independent experiments (*n* = 3) with 20-25 GUVs analyzed per experiment.

## References

1. Bozelli, J.C., Jr. and R.M. Epand, Membrane Shape and the Regulation of Biological Processes. J Mol Biol, 2020. 432(18): p. 5124–5136.

2. Daumke, O., A. Roux, and V. Haucke, BAR domain scaffolds in dynamin-mediated membrane fission. Cell, 2014. 156(5): p. 882–92.

3. Haucke, V. and M.M. Kozlov, *Membrane remodeling in clathrin-mediated* endocytosis. J Cell Sci, 2018. 131(17).

4. McMahon, H.T. and E. Boucrot, Molecular mechanism and physiological functions of clathrin-mediated endocytosis. Nat Rev Mol Cell Biol, 2011. 12(8): p. 517–33.

5. McMahon, H.T. and J.L. Gallop, *Membrane curvature and mechanisms of* dynamic *cell membrane remodelling*. Nature, 2005. 438(7068): p. 590–6.

6. Kozlov, M.M. and J.W. Taraska, *Generation of nanoscopic membrane curvature* for *membrane trafficking*. Nat Rev Mol Cell Biol, 2023. 24(1): p. 63–78.

7. Lundmark, R. and S.R. Carlsson, Driving membrane curvature in clathrin-dependent and clathrin-independent endocytosis. Semin Cell Dev Biol, 2010. 21(4): p. 363–70.

8. Boucrot, E., et al., Endophilin marks and controls a clathrin-independent endocytic pathway. Nature, 2015. 517(7535): p. 460–5.

9. Casamento, A. and E. Boucrot, Molecular mechanism of Fast Endophilin-Mediated Endocytosis. Biochem J, 2020. 477(12): p. 2327–2345.

10. Watanabe, S. and E. Boucrot, Fast and ultrafast endocytosis. Curr Opin Cell Biol, 2017. 47: p. 64–71.

11. Genet, G., et al., *Endophilin-A2 dependent VEGFR2 endocytosis promotes* sprouting *angiogenesis*. Nat Commun, 2019. 10(1): p. 2350.

12. Bertot, L., et al., Quantitative and Statistical Study of the Dynamics of Clathrin-Dependent and -Independent Endocytosis Reveal a Differential Role of EndophilinA2. Cell Reports, 2018. 22(6): p. 1574–1588.

13. Renard, H.F., et al., *Endophilin-A2 functions in membrane scission in clathrin-*independent *endocytosis*. Nature, 2015. 517(7535): p. 493–6.

14. Renard, H.F., et al., *Endophilin-A3 and Galectin-8 control the clathrin-*independent *endocytosis of CD1CC*. Nat Commun, 2020. 11(1): p. 1457.

15. Gallop, J.L., et al., Mechanism of endophilin N-BAR domain-mediated membrane curvature. EMBO Journal, 2006. 25(12): p. 2898–2910.

16. Mim, C., et al., Structural basis of membrane bending by the N-BAR protein endophilin. Cell, 2012. 149(1): p. 137–45.

17. Li, P., et al., Phase transitions in the assembly of multivalent signalling proteins. Nature, 2012. 483(7389): p. 336–340.

18. Day, K.J., et al., *Liquid-like protein interactions catalyse assembly of endocytic* vesicles. Nature Cell Biology, 2021. 23(4): p. 366–376.

19. Kozak, M. and M. Kaksonen, Condensation of Ede1 promotes the initiation of endocytosis. eLife, 2022. 11: p. e72865.

20. Mamta, M., et al., Adaptor interactions trigger Pan1 self-assembly during clathrin mediated endocytosis in budding yeast. bioRxiv, 2026.

21. Malady, B., et al., Endocytic accessory proteins assemble clathrin while simultaneously destabilizing protein condensates. bioRxiv, 2026.

22. Sun, Y., et al., Phosphoregulation of hierarchical protein condensate formation during clathrin-mediated endocytosis. bioRxiv, 2026.

23. Banjade, S. and M.K. Rosen, Phase transitions of multivalent proteins can promote clustering of membrane receptors. eLife, 2014. 3: p. e04123.

24. Su, X., et al., Phase separation of signaling molecules promotes T cell receptor signal transduction. Science, 2016. 352(6285): p. 595–599.

25. Mondal, S., et al., Multivalent interactions between molecular components involved in fast endophilin mediated endocytosis drive protein phase separation. Nature Communications, 2022. 13(1): p. 5017.

26. Narayan, K.B., et al., Endophilin-lamellipodin-VASP, key components in fast endophilin-mediated endocytosis, control actin polymerization within liquid-like condensates. Journal of Biological Chemistry, 2025. 301(12): p. 110834.

27. Boucrot, E., et al., Endophilin marks and controls a clathrin-independent endocytic pathway. Nature, 2015. 517(7535): p. 460–465.

28. Chan Wah Hak, L., et al., FBP17 and CIP4 recruit SHIP2 and lamellipodin to prime the plasma membrane for fast endophilin-mediated endocytosis. Nature Cell Biology, 2018. 20(9): p. 1023–1031.

29. Farsad, K., et al., Generation of high curvature membranes mediated by direct endophilin bilayer interactions. Journal of Cell Biology, 2001. 155(2): p. 193–200.

30. Gowrisankaran, S., et al., Endophilin-A coordinates priming and fusion of neurosecretory vesicles via intersectin. Nature Communications, 2020. 11(1): p. 1266.

31. Sundborger, A., et al., An endophilin-dynamin complex promotes budding of clathrin-coated vesicles during synaptic vesicle recycling. Journal of Cell Science, 2011. 124(1): p. 133–143.

32. Capraro, B.R., et al., Kinetics of endophilin N-BAR domain dimerization and membrane interactions. Journal of Biological Chemistry, 2013. 288(18): p. 12533–12543.

33. Angelova, M.I. and D.S. Dimitrov, Liposome electroformation. Faraday Discussions of the Chemical Society, 1986. 81(0): p. 303–311.

34. Kenausis, G.L., et al., Poly(l-lysine)-g-Poly(ethylene glycol) Layers on Metal Oxide Surfaces: Attachment Mechanism and Effects of Polymer Architecture on Resistance to Protein Adsorption. The Journal of Physical Chemistry B, 2000. 104(14): p. 3298–3309.

35. Melo, A.M., S. Elbaum-Garfinkle, and E. Rhoades, Insights into tau function and dysfunction through single-molecule fluorescence. Methods in Cell Biology, 2017. 141: p. 27–44.

36. Vehlow, A., et al., Endophilin, lamellipodin, and mena cooperate to regulate F-actin-dependent EGF-receptor endocytosis. EMBO Journal, 2013. 32(20): p. 2722–2734.

37. Hansen, S.D. and R.D. Mullins, Lamellipodin promotes actin assembly by clustering Ena/VASP proteins and tethering them to actin filaments. eLife, 2015. 4(AUGUST2015).

38. Huttner, W.B. and A. Schmidt, Lipids, lipid modification and lipid–protein interaction in membrane budding and fission — insights from the roles of endophilin A1 and synaptophysin in synaptic vesicle endocytosis. Current Opinion in Neurobiology, 2000. 10(5): p. 543–551.

39. Weissenhorn, W., Crystal Structure of the Endophilin-A1 BAR Domain. Journal of Molecular Biology, 2005. 351(3): p. 653–661.

40. Chen, Z., et al., The N-Terminal Amphipathic Helix of Endophilin Does Not Contribute to Its Molecular Curvature Generation Capacity. Journal of the American Chemical Society, 2016. 138(44): p. 14616–14622.

41. Narayan, K.B., et al., Endophilin-Lamellipodin-VASP, key components in fast endophilin-mediated endocytosis, control actin polymerization within liquid-like condensates. bioRxiv, 2024.

42. Masuda, M., et al., Endophilin BAR domain drives membrane curvature by two newly identified structure-based mechanisms. The EMBO Journal, 2006. 25(12): p. 2889.

43. Yang, L.Q., A.F. Huang, and W.D. Xu, Biology of endophilin and its role in disease. Frontiers in Immunology, 2023. 14: p. 1297506.

44. Kjaerulff, O., L. Brodin, and A. Jung, The Structure and Function of Endophilin Proteins. Cell Biochemistry and Biophysics 2010 60:3, 2010. 60(3): p. 137-154.

45. Cestra, G., et al., The SH3 domains of endophilin and amphiphysin bind to the proline-rich region of synaptojanin 1 at distinct sites that display an unconventional binding specificity. Journal of Biological Chemistry, 1999. 274(45): p. 32001–32007.

46. De Heuvel, E., et al., Identification of the major synaptojanin-binding proteins in brain. Journal of Biological Chemistry, 1997. 272(13): p. 8710–8716.

47. Ringstad, N., Y. Nemoto, and P. De Camilli, Differential Expression of Endophilin 1 and 2 Dimers at Central Nervous System Synapses. Journal of Biological Chemistry, 2001. 276(44): p. 40424–40430.

48. Sundborger, A., et al., An endophilin–dynamin complex promotes budding of clathrin-coated vesicles during synaptic vesicle recycling. Journal of Cell Science, 2011. 124(1): p. 133–143.

49. Harmon, T.S., et al., Intrinsically disordered linkers determine the interplay between phase separation and gelation in multivalent proteins. Elife, 2017. 6.

